# Phosphatase specificity influences phosphorylation timing of CDK substrates during the cell cycle

**DOI:** 10.1101/2025.02.21.639482

**Authors:** Theresa U. Zeisner, Tania Auchynnikava, Paul Nurse

## Abstract

Cell cycle events are ordered by cyclin-dependent kinases (CDKs), which phosphorylate hundreds of substrates. Multiple phosphatases oppose CDK substrate phosphorylation, yet a systematic understanding of how these phosphatases collectively influence phosphorylation timing is lacking. Here, we show that phosphatases influence the timing of CDK substrate phosphorylation during G2 and mitosis in fission yeast. We identify substrates of four phosphatases (PP2A-B55, PP2A-B56, CDC14, and PP1), showing that each phosphatase targets a distinct subset of CDK substrate sites. On average, sites dephosphorylated by CDC14 and PP2A-B56 are phosphorylated earlier during G2, followed by sites dephosphorylated by PP1 and then PP2A-B55. This suggests that the identity of the phosphatase impacts the timing of CDK substrate phosphorylation, establishing different phosphorylation thresholds at the G2/M transition. Consistent with this, depletion of PP2A-B55 and CDC14 advances mitotic onset independently of CDK activity regulation, likely due to the earlier phosphorylation of their respective CDK substrates.

## Introduction

The events of the cell cycle are driven by the reversible phosphorylation of hundreds of phosphosites on CDK substrates (hereafter referred to as CDK substrate sites). These phosphorylations must occur in a distinct temporal order to ensure proper cell cycle progression and, ultimately, the production of viable daughter cells^1^. This phosphorylation order is achieved by the *in vivo* sensitivity of CDK substrate sites to increasing CDK-to-phosphatase activity^2,3^. CDK activity is low during G1 and S-phase, resulting in the phosphorylation of S-phase but not mitotic substrates^2^. Dephosphorylation of CDK’s Tyrosine15 residue at the G2/M transition leads to a switch-like increase in CDK activity and the phosphorylation of CDK substrates at mitosis^4,5^. However, increasing CDK activity alone does not order CDK substrate phosphorylations within the cell cycle, since the efficiency with which a CDK substrate is phosphorylated by CDK *in vitro* does not correlate with the timing of its phosphorylation during the cell cycle^6^. Thus, *in vivo,* additional factors, such as opposing phosphatase activity, must impact the timing of CDK substrate phosphorylation. Multiple phosphatases have been reported to target CDK substrates^7,8^, but a systematic understanding of how these phosphatases together shape the phosphorylation timing of CDK substrates during interphase is currently lacking.

So far, CDK-opposing phosphatases, including PP2A, PP1, and in yeast also CDC14, have been studied in-depth for their crucial roles during mitosis and at mitotic exit through dephosphorylating CDK substrates^9–13^. However, these phosphatases are also active during interphase and can continuously remove phosphate groups from CDK substrate sites. Consequently, on the level of individual molecules, phosphorylation sites undergo repeated cycles of phosphorylation and dephosphorylation. The overall amount of phosphorylated substrate (hereafter referred to as net phosphorylation) is therefore determined by the catalytic activities of both the kinase and the opposing phosphatase. This turnover of phosphate groups is readily detected when CDK activity is chemically inhibited because CDK substrate sites become rapidly dephosphorylated, with half-lives of ∼2 minutes^2,14^. A more rapid phosphatase-dependent turnover of a CDK substrate site would set a higher threshold that CDK activity must achieve to bring about net phosphorylation of this site, thus delaying net phosphorylation until CDK activity is higher^15^. Consistent with this, inhibition of PP2A-B55^CDC55^ in budding yeast advances the phosphorylation of threonine sites and the CDK substrate Ndd1^16^. However, how other CDK-opposing phosphatases affect the timing of substrate phosphorylation during interphase and how they interact together remains largely unknown. Genetic screens, and biochemical and cell-based assays suggest that PP2A-B55, PP2A-B56, PP1 and CDC14 are important cell cycle regulators, affecting processes such as chromosome biorientation, the spindle assembly checkpoint, and mitotic entry in a range of organisms^7,8^. Despite the importance of these phosphatases for various cell cycle events, we do not know which phosphatases target the majority of CDK substrate sites. This may be partly due to the historical misconception that phosphatases are unspecific housekeeping enzymes^17^.

In this paper, we investigate the role of CDK-opposing phosphatases in regulating the timing of CDK substrate phosphorylation and, thus, temporal ordering within the cell cycle in fission yeast. Using a systematic phosphoproteomic approach, we identified *in vivo* substrates of four CDK-opposing phosphatases: PP2A-B55, PP2A-B56, PP1 and CDC14. Our data show that these four phosphatases target distinct subgroups of CDK substrate sites, which are, on average, net phosphorylated at different times during the cell cycle. This suggests that the division of labour between the phosphatases impacts the timing of CDK substrate phosphorylation during G2 onto the onset of mitosis.

## Results

### Identification of CDK-dependent phosphatase substrates *in vivo*

We identified which CDK substrate sites are targeted by PP2A-B55, PP2A-B56, PP1, and CDC14 *in vivo* using phosphoproteomics by comparing the dephosphorylation kinetics of CDK substrate sites in the presence and absence of each phosphatase. To identify CDK substrate sites, we synchronised cells in G2 with an ATP analogue-sensitive CDK allele (*cdc2-as*) and released them into mitosis, at which point CDK substrate sites are highly phosphorylated. The dephosphorylation of phosphosites upon CDK inhibition with the ATP analogue 1-NmPP1 was monitored using phosphoproteomics (Fig. 1a and Extended Data Fig. 1a,b). We classified phosphosites as CDK substrate sites if they were dephosphorylated rapidly upon CDK inhibition by fitting an exponential decay (Extended Data Fig. 1c-e). The identified CDK substrate sites were enriched for the full CDK consensus motif ([ST]Px[KR]) and for gene ontology terms associated with the cell cycle^18^ (Extended Data Fig. 1f,g). Consistent with previous data, CDK substrate sites had a median half-life of <2 mins, whilst protein levels remained stable (Extended Data Fig. 1h,i)^2^. Around 15 % of CDK substrate sites had a half-life of less than 30 seconds, emphasising the rapid turnover of these phosphosites (Supplementary Table 1). The dephosphorylation rates of CDK substrate sites were similar between four phosphoproteomic experiments, indicating reproducibility between biological repeats (Extended Data Fig. 1j).

**Figure 1:**
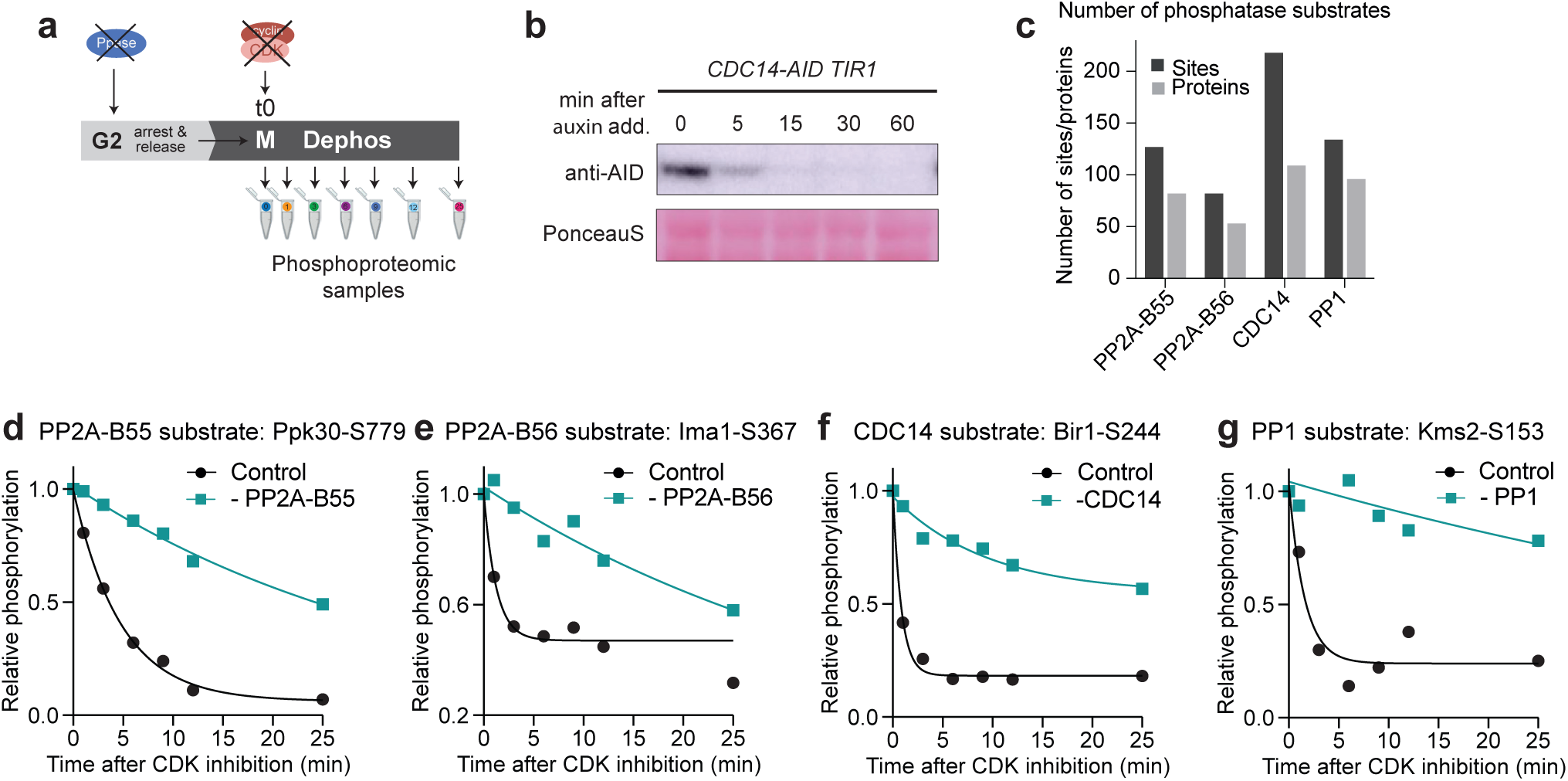
Identification of *in vivo* CDK-dependent phosphatase substrates. **a)** Schematic of the experimental set-up to identify phosphatase substrates in a synchronised culture upon CDK inhibition. The AID system was used to degrade the phosphatases of interest, the ATP-analogue 1-NmPP1 was used to inhibit CDK activity in an analogue-sensitive strain. **b)** Western blot against the AID-tag of CDC14, showing degradation upon auxin addition. Ponceau-S stain was used for total protein normalisation. **c)** Grouped bar graphs showing the number of CDK-dependent phosphatase substrate sites (dark grey) and proteins (light grey). **d-g)** Examples of identified phosphatase substrates. Relative phosphorylation level in the presence (black) and absence (turquoise) of phosphatase of interest upon CDK inhibition. Curves are a one-phase exponential decay fitted to the relative phosphorylation level determined by phosphoproteomics.

The rapid dephosphorylation of CDK substrate sites indicates that phosphatases are active. We, therefore, expected that in the absence of the opposing phosphatase, the dephosphorylation of a site to be slower. Thus, to identify which phosphatases dephosphorylate which specific CDK substrate sites, we quantified the dephosphorylation of CDK substrate sites in the absence of each of the four phosphatases. Using an auxin-inducible degron (AID) system, we individually depleted the activity of the four CDK-opposing phosphatases^19,20^. PP2A and PP1 are both holoenzymes consisting of different subunits (Extended Data Fig. 2a). For PP2A, we individually degraded the regulatory subunits B55 (encoded by *pab1*) and B56 (encoded by *par1* and *par2*), thus abolishing the activity of two specific PP2A holoenzymes separately. PP1 associates with multiple regulatory subunits; for example, there are >200 regulatory PP1 subunits in human cells^21^. We, therefore, degraded both catalytic subunits of PP1 together (encoded by *dis2* and *sds21*), thereby inhibiting the activity of all possible PP1 holoenzymes. We also degraded CDC14 (encoded by *clp1*), which does not form a holoenzyme. Degradation of the phosphatases was rapid, occurring within 15 minutes and led to phenotypes similar to previously reported deletion phenotypes of the corresponding genes (Fig. 1b and Extended Data Fig. 2b-d).

We classified CDK substrate sites as phosphatase targets if their dephosphorylation was substantially slower in the absence of one of the phosphatases by comparing the dephosphorylation in the control and phosphatase-degraded condition (Extended Data Fig. 1c and Extended Data Fig. 3a). We identified between 80-220 CDK-dependent phosphatase substrate sites for each of the four investigated phosphatases, together totalling 561 sites on 262 proteins (Fig. 1c-g and Extended Data Fig. 3b and Supplementary Table 1). For each of the investigated phosphatases, our dataset included phosphatase substrates that have been previously identified, such as the PP2A-B55 substrate PRC1 and the CDC14 substrate MSH6^22,23^ (Supplementary Table 1). Overall, this approach provides a comprehensive dataset of CDK-dependent and, thus, cell cycle-relevant phosphatase substrates.

### Phosphatases target distinct subgroups of CDK substrate sites

Our dataset enabled us to determine if one or multiple phosphatases dephosphorylated a specific CDK substrate site (Fig. 2a and Extended Data Fig. 4a-c). To allow comparisons across all datasets, only sites detected in all four experiments were used for this analysis (274 sites) (Extended Data Fig. 4d,e). We identified the opposing phosphatase for 59 % of these common CDK substrate sites (Fig. 2b). PP2A-B55 and CDC14 both targeted around 20 % of CDK substrate sites, while PP1 and PP2A-B56 targeted 11 % and 2 %, respectively. There was very little overlap between the groups of phosphatase substrates, with less than 5 % of CDK substrate sites being dephosphorylated by multiple phosphatases (Fig. 2b and Extended Data Fig. 4f,g). This indicates that these phosphatases target distinct substrate pools *in vivo*, countering the notion that phosphatases are unspecific. This division of labour between phosphatases in opposing CDK substrate sites raises two questions: what is the molecular basis for this apparent substrate specificity, and what regulatory consequences does this division of labour have for the timing of CDK substrate phosphorylation.

**Figure 2:**
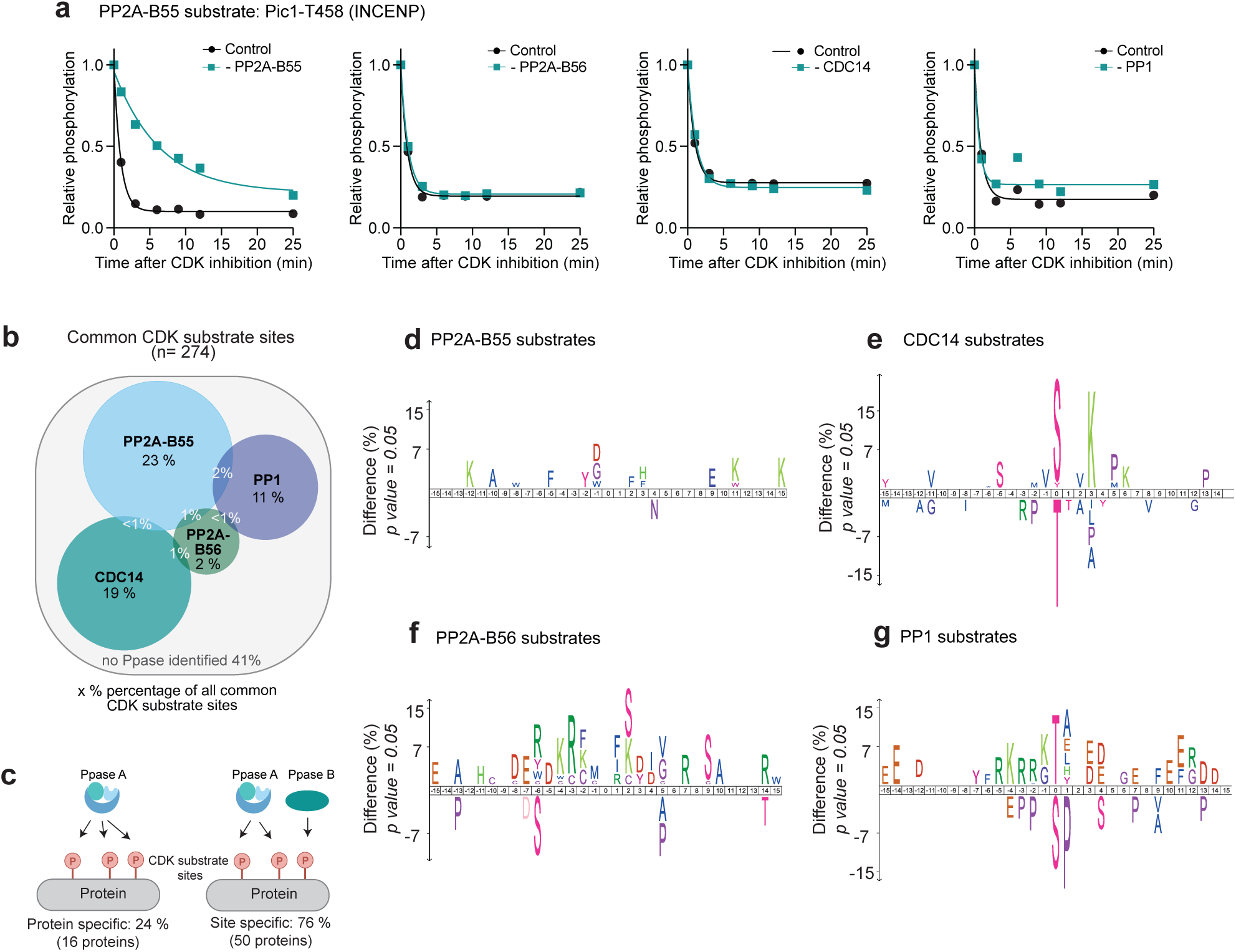
Phosphatases target distinct subgroups of CDK substrate sites. **a)** Example of a specific PP2A-B55 phosphatase substrate. Relative phosphorylation level of the INCENP orthologue Pic1-T458 in the presence (black) and absence (turquoise) of the indicated phosphatase upon CDK inhibition. Pic1-T458 dephosphorylation is substantially slowed in the absence of PP2A-B55 (left panel), but not in the absence of PP2A-B56, CDC14 or PP1.Curves are a one-phase exponential decay fitted to the relative phosphorylation level determined by phosphoproteomics. **b)** Venn diagram depicting the overlap of CDK-dependent phosphatase substrates of PP2A-B55, PP2A-B56, PP1 and CDC14 (using only CDK-dependent sites common to all four experimental sets as background). **c)** Schematic of protein-specific or site-specific phosphatase substrates on proteins with multiple CDK-dependent sites. **d-g)** Amino acid motifs of CDK-dependent phosphatase substrates of the four different phosphatases. A 31 amino acid sequence centred on the phosphorylated residue of the identified phosphatase substrates was compared against all identified CDK substrate sites using iceLogo (p-value=0.05)^65^. Amino acids above the line are enriched, while those below the line are depleted in the phosphatase substrates.

We first assessed whether phosphatase specificity arises at the protein or phosphosite level, by examining proteins with multiple CDK substrate sites. For 76 % of multiple-phosphorylated CDK substrate proteins, different phosphosites on the same protein were targeted by different phosphatases (Fig. 2c). This indicates that phosphatase substrate specificity operates at the level of individual phosphosites rather than the level of the entire protein. To interrogate the molecular basis of this, we investigated whether sites targeted by a given phosphatase shared common features. Comparing the sequences of the phosphatase substrate sites with that of all CDK substrate sites revealed differences in the amino acids surrounding phosphorylation sites (Fig. 2d-g). PP2A-B56 substrates were enriched for basic amino acids surrounding the phosphorylation site (Fig. 2f), and CDC14 substrates were enriched for phosphoSerine (pSerines) and lysine at the +3 position, which corresponds to the full CDK consensus motif^18^ (Fig. 2e). PP1 substrates were strongly enriched for phosphoThreonine (pThreonines) at the phosphorylation site (Fig. 2g), even though around half of CDK-dependent PP1 substrates are pSerines. This can be explained by the higher abundance of serines than threonines in the phosphoproteome^24^ (Extended Data Fig. 4h). This result emphasises the need to identify individual substrates of phosphatases experimentally under *in vivo* conditions, rather than relying solely on consensus motifs for their identification. Consistent with previous work, PP2A-B55 and PP1 dephosphorylated pThreonine CDK substrate sites significantly faster than pSerine sites^25–27^ (Extended Data Fig. 4i-l).

We conclude that CDK-dependent phosphatase substrates are site-specific rather than protein-specific, and site-specificity is at least partly determined by the different amino acid preferences of each phosphatase.

### Identification of CDK-independent phosphatase substrates reveals conservation of phosphatase substrate motifs

The investigated phosphatases are also known to dephosphorylate sites, not phosphorylated by CDK and we therefore extended our analysis to identify which non-CDK substrates are targeted by the four phosphatases^8^. We reasoned that the phosphorylation levels of sites phosphorylated by kinases other than CDK would remain unchanged immediately after inhibiting CDK activity in mitosis. We used the same phosphoproteomic dataset to identify sites where the phosphorylation level remained unchanged upon CDK inhibition, classifying them as CDK-independent sites (Fig. 3a and Extended Data Fig. 5a,b). CDK-independent sites for which the phosphorylation level was significantly higher in the absence of the phosphatase were classified as CDK-independent phosphatase substrates (Fig. 3a). This analysis identified between 600-1,300 phosphatase substrates for each of the four investigated phosphatases, totalling over 3,600 sites (Extended Data Fig. 5c,d, Supplementary Table 2). Combining the CDK-dependent and CDK-independent phosphatase substrates together gave an estimate of the total substrate sites targeted by each phosphatase (Fig. 3b). As before, to compare between datasets, we only included sites detected in all four experimental sets for this analysis. Similar to the CDK-dependent phosphatase substrates, there was limited overlap between the different groups of total phosphatase substrates, with less than 3 % of sites targeted by multiple phosphatases (Fig. 3b). We estimated that PP1 and PP2A-B55 each targeted approximately 6-9 % of all phosphosites in the phosphoproteome, while PP2A-B56 and CDC14 each targeted an additional 3-4 % of all phosphosites, bringing the total to around 25 % of the entire phosphoproteome (Fig. 3b).

**Figure 3:**
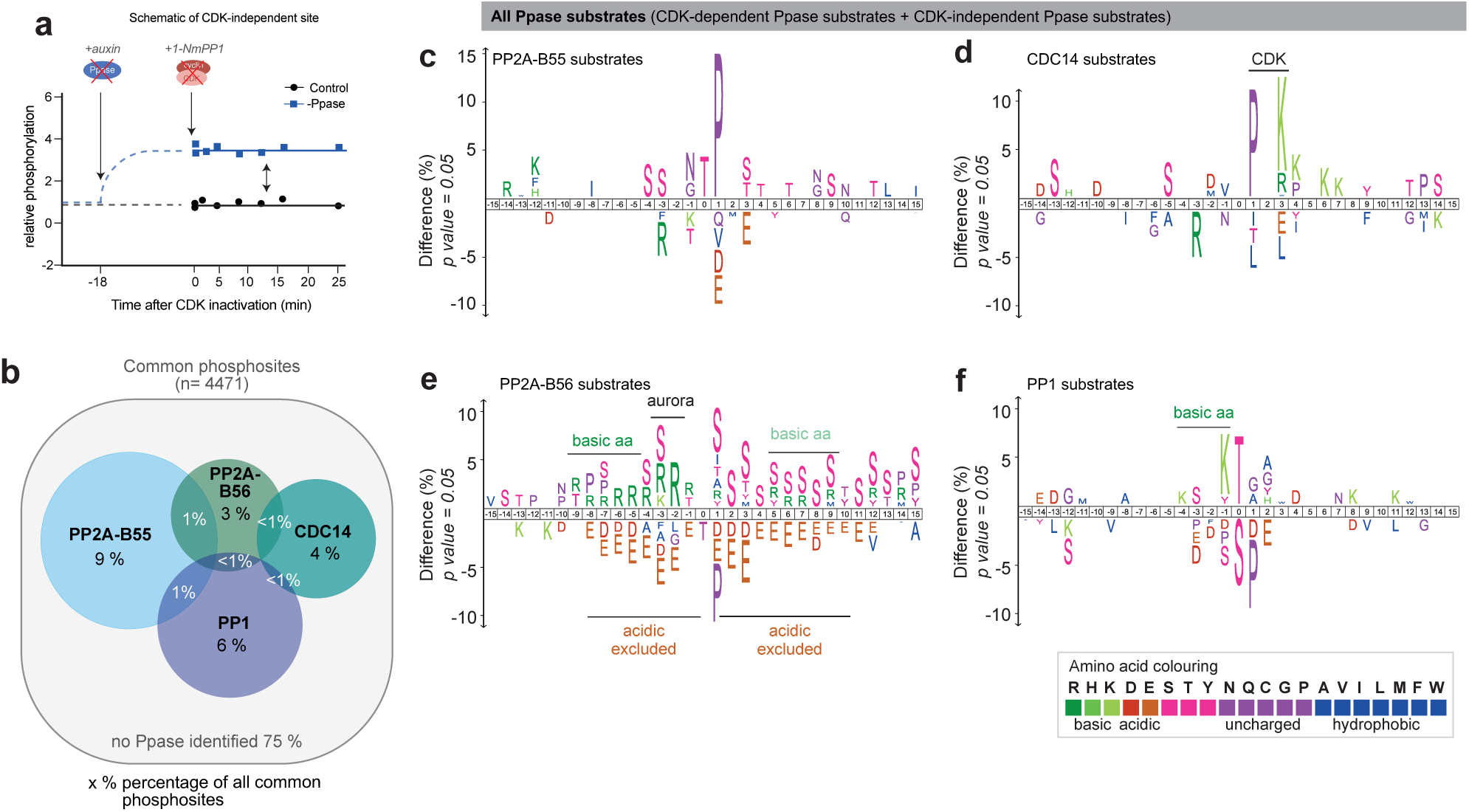
Phosphatases target distinct subgroups of CDK-independent sites. **a)** Schematic of the relative phosphorylation level of a CDK-independent phosphatase substrate in the presence (black) and absence (blue) of the phosphatase. **b)** Venn diagram depicting the overlap of total phosphatase substrates, using only phosphosites common to all four experimental sets. **c-f)** Amino acid motifs of total phosphatase substrates (CDK-dependent + CDK-independent phosphatase substrates) of the four investigated phosphatases. A 31 amino acid sequence centred on the phosphorylated residue of the identified phosphatase substrates was compared against all identified phosphosites sites using iceLogo (p-value=0.05)^65^. Amino acids above the line are enriched, while those below the line are depleted in the phosphatase substrates.

Taking the CDK-dependent and CDK-independent phosphatase substrates together revealed distinct motifs of amino acids surrounding the phosphorylated residue for PP2A-B55, PP2A-B56, PP1, and CDC14 substrates (Fig. 3c-f). PP2A-B55 substrates were enriched for pThreonine followed by a Proline, while these residues were depleted in PP2A-B56 substrates (Fig. 3c,e). PP2A-B56 substrates were strongly enriched in basic amino acids at the -2 and -3 position, which resembles the Aurora B kinase motif^28^. CDC14 substrates were enriched for PxK (+1 to +3 position), consistent with the full CDK consensus motif^18^. PP1 substrates were enriched for pThreonines, basic amino acids upstream of phosphorylation sites, and depleted for prolines at the +1 position. CDK also phosphorylates non-proline sites (non-canonical CDK substrate sites)^29^ and consistent with this, approximately 50 % of CDK-dependent PP1 substrates were non-canonical CDK sites (Supplementary Table 1). Overall, the phosphatase motifs showed a high similarity with motifs of metazoan phosphatases^22,26,27,30^, indicating conservation of substrate specificity across eukaryotes.

To investigate which kinases these phosphatases oppose in addition to CDK, we conducted an enrichment analysis based on the kinase substrate motifs from the human kinome atlas^31^. This revealed that PP2A-B55 substrates were enriched for MAPK motifs and, as expected, CDK kinase motifs, while PP2A-B56 substrates were strongly enriched for Aurora B kinase motifs^28^ (Extended Data Fig. 5e). Additionally, we also searched the groups of total phosphatase substrates for consensus motifs of cell cycle-relevant kinases, including CDK, Aurora B, Polo, NIMA, NDR2 and DDK^18,28,32–36^. This confirmed the preference of PP2A-B56 for Aurora B substrates and of PP2A-B55 and CDC14 for CDK substrates. It further emphasised that each of the phosphatases targets a range of cell-cycle kinase substrates (Extended Data Fig. 5f,g). In addition to preferences for amino acids surrounding the phosphorylated residue, phosphatases can additionally enhance substrate specificity by binding to short linear motifs (SLiMs)^37^. PP2A-B56 substrates were enriched for LxxIxE motifs known to interact with a conserved region on B56, while CDC14 substrates were enriched for PxL motifs (Extended Data Fig. 5h)^38,39^.

This analysis revealed that the phosphatases target substrates of multiple cell cycle protein kinases, including Polo and Aurora B, and emphasises that kinases and phosphatases generally do not act in pairs; rather, a single phosphatase targets substrates from several different kinases. Together, these data show that the investigated phosphatases are specific enzymes which target distinct groups of substrates *in vivo*.

### CDK substrates opposed by different phosphatases are net phosphorylated at different cell cycle times

We next investigated whether the division of labour between phosphatases in opposing distinct subgroups of CDK substrate sites had a functional impact on the timing of CDK substrate phosphorylation. In particular, we analysed a phosphoproteomic timecourse dataset to determine whether the net phosphorylation timing of CDK substrates differed between the groups of phosphatase substrates. This dataset quantified CDK substrate phosphorylations during a synchronised cell cycle following release from a G2 arrest in the presence of all phosphatases^2^. Comparing the median phosphorylation profiles of CDK substrate sites dephosphorylated by different phosphatases revealed a temporal order during the cell cycle (Fig. 4a). CDK substrate sites targeted by PP2A-B56 and CDC14 were, on average, net phosphorylated earlier during the cell cycle, followed by CDK substrate sites targeted by PP1 and then PP2A-B55. The median net phosphorylation of CDK/CDC14 sites started to increase around 20 minutes earlier than CDK/PP2A-B55 sites (Fig. 4a).

**Figure 4:**
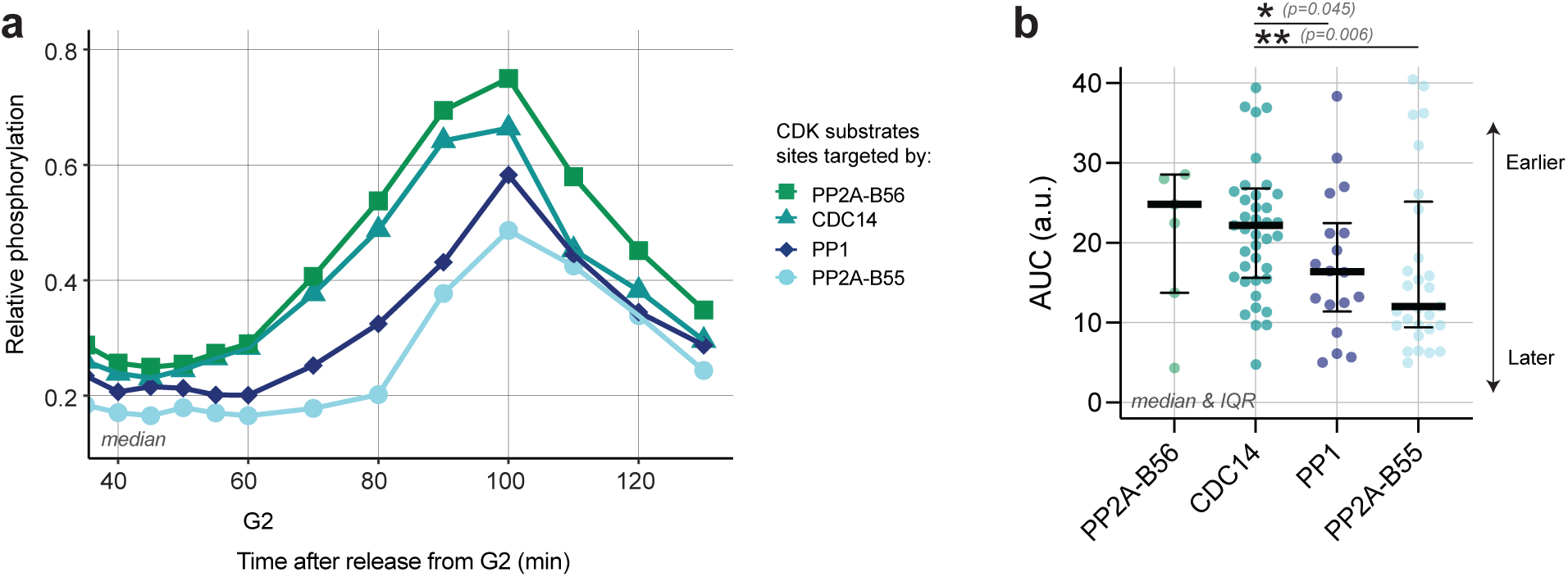
Net phosphorylation timing of CDK substrate sites differs between phosphatase substrate groups. **a)** Median relative phosphorylation of CDK-dependent phosphatase substrates during a synchronised cell cycle, identified by mass spectrometry (n-numbers: PP2A-B56:7, CDC14: 41, PP1: 18 and PP2A-B55=29). **b)** Beeswarm plot of AUC values of CDK-dependent phosphatase substrate sites, split according to which phosphatase dephosphorylates the CDK substrate site (CDC14, PP2A-B56, PP1 and PP2A-B55). AUC values are the integral of the relative phosphorylation (50-100 mins after release from G2) and act as a proxy for phosphorylation timing and pattern during the cell cycle. Error bars correspond to median and inter-quartile range (IQR). Significant differences were determined using the Mann-Whitney test (two-tailed), only significant differences between groups are shown on graph.

To quantify the phosphorylation timing of individual CDK substrate sites, we calculated the area under the curve (AUC) of the phosphorylation level for each CDK substrate from G2 phase to mitosis (50-100 mins after release from a G2 arrest). The later a substrate is phosphorylated, the smaller is its AUC (Extended Data Fig. 6a). This analysis revealed a spread of phosphorylation timings within each group of phosphatase substrates (Fig. 4b). The median AUC values confirm the earlier finding that, on average, CDK substrate sites targeted by PP2A-B56 and CDC14 were net phosphorylated earlier (higher median AUC), while sites targeted by PP2A-B55 and PP1 were net phosphorylated significantly later (lower median AUC). It also highlighted that S-phase substrates are not all targeted by the same phosphatase. We conclude that the identity of the phosphatase that dephosphorylates a CDK substrate appears to impact the timing of CDK substrate phosphorylation during G2 and mitosis but does not distinguish between S-phase and mitotic CDK substrates.

There is a considerable spread in the timing of CDK substrate phosphorylations within each group of phosphatase substrates (Fig. 4b). This indicates that the exact phosphorylation timing of each CDK substrate site can be expected to be the consequence of several factors, including how well a site is phosphorylated by CDK, the identity of the opposing phosphatase, the presence of SLiMs, and the sub-cellular localisation of the substrate^15,40,41^. To stratify the contributions of some of these factors, we grouped CDK substrates according to these criteria (Extended Data Fig. 6b). By comparing the difference of median AUCs between sub-categories of each of these factors, we can estimate the size of their effects. This showed that the identity of the phosphatase and substrate localisation had the greatest difference in AUC values between groups, being 4 and 3 times larger compared to AUC differences seen when CDK substrates are grouped according to the phosphorylated residue (Extended Data Fig. 6b). This indicates that both the identity of the phosphatase and sub-cellular localisation of the CDK substrates have strong effects on the cell cycle timing of CDK substrate phosphorylation. The observed differences in the phosphorylation timing of CDK substrates, categorised by phosphatases or subcellular location, are independent. If these two categories were connected, CDC14 and PP2A-B56 substrates, which are phosphorylated first during the cell cycle, should be predominantly located in the nucleus, as that is the compartment in which CDK activity rises first^42^. However, none of the phosphatase substrate groups were significantly enriched in any specific subcellular compartment (Extended Data Fig. 6c). Furthermore, if we restrict sites to a specific sub-cellular compartment, such as the nucleus or cytoplasm, CDC14 substrate sites were still phosphorylated before PP2A-B55 sites, regardless of the cellular compartment (Extended Data Fig. 6d). This reinforces the notion that the differences in phosphorylation timing of CDK substrates categorised by phosphatases or subcellular localisation act independently of one another.

We conclude that the division of labour between the phosphatases that dephosphorylate distinct subgroups of CDK substrate sites plays a role in ordering the phosphorylation of CDK substrate sites during the transition from G2 into mitosis. Our data showed that CDK substrate sites opposed by CDC14 were, on average, phosphorylated earlier than those opposed by PP1 and PP2A-B55.

### PP2A-B55 impacts the phosphorylation timing of a CDK/PP2A-B55 substrate *in vivo*

Apart from the identity of the phosphatase, how efficiently a CDK substrate is dephosphorylated by a given phosphatase may also affect phosphorylation timing within the cell cycle^15^. Since CDK substrate sites targeted by PP2A-B55 were, on average, phosphorylated significantly later than the average CDK substrate site during G2, we interrogated the possibility that PP2A-B55 may delay net phosphorylation of its CDK substrate sites. To do so, we used an *in vivo* phosphorylation-sensor to determine whether a PP2A-B55 site is net phosphorylated earlier during the cell cycle in the absence of the phosphatase. We identified Cut3-T19, which is part of the *in vivo* phospho-sensor SynCut3-mC, as a PP2A-B55 substrate site (Fig. 5a,b). Upon net phosphorylation of T19 by CDK, Cut3 translocates from the cytoplasm into the nucleus at the onset of mitosis^43^, so the nuclear to cytoplasmic ratio of exogenously expressed SynCut3-mC can be used to monitor its phosphorylation status (Fig. 5c). In the presence of PP2A-B55 the net phosphorylation of T19 is low during interphase and abruptly increases ∼ 12 minutes before peak mitosis. In the absence of PP2A-B55 the net phosphorylation of T19 begins earlier in G2, on average 28 mins before the mitotic peak, and its initial net phosphorylation increases less rapidly (Fig. 5d,e and Extended Data Fig. 7a). This change in SynCut3-T19 phosphorylation pattern is not observed in the absence of other cell cycle phosphatases (CDC14, Pyp1, PP1 catalytic subunit), indicating that this is a phosphatase-specific effect (Extended Data Fig. 7b,c).

**Figure 5:**
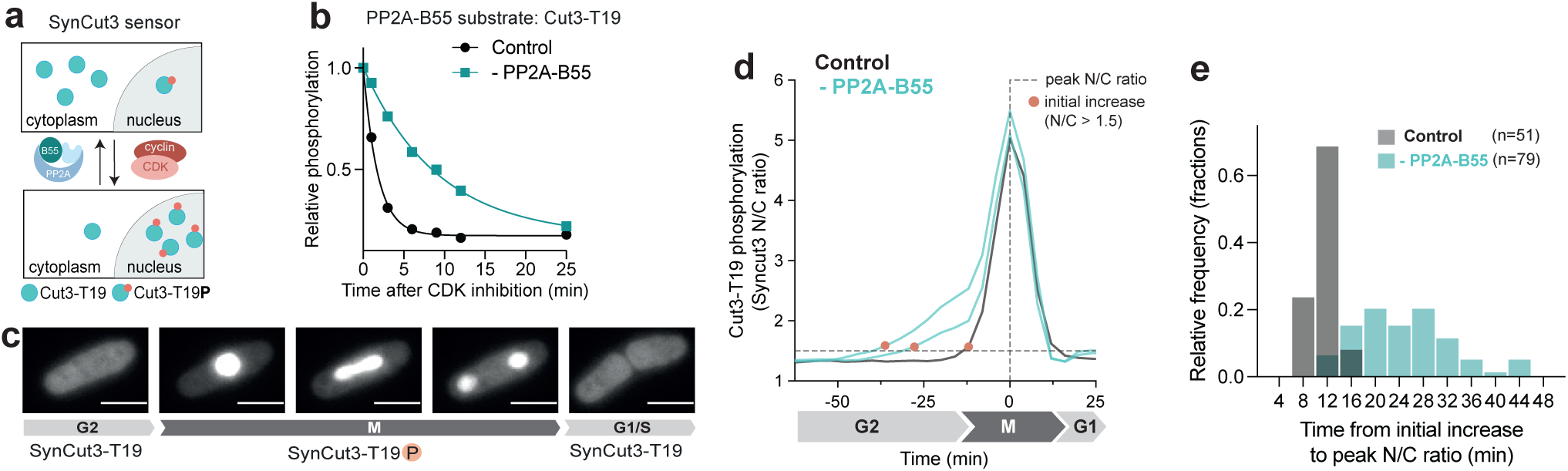
In the absence of PP2A-B55 the CDK substrate site Cut3-T19 is phosphorylated earlier and less rapidly. **a)** Schematic of SynCut3-mCherry sensor. **b)** Relative phosphorylation level of Cut3-T19 upon CDK-inhibition in the presence (black) and absence (turquoise) of PP2A-B55. Curves are a one-phase exponential decay fitted to the relative phosphorylation level determined by phosphoproteomics. **c)** Representative fluorescent images of SynCut3-mCherry throughout a cell cycle within a single fission yeast cell, nuclear import of the sensor and binucleation can be seen in image 2-4. Error bars correspond to 5 µm. **d)** Representative single-cell traces of SynCut3-mC N/C ratio in the presence (black) and absence (turquoise) of PP2A-B55, as determined by life-cell fluorescent imaging. Traces are aligned at their peak N/C ratio. The initial increase in N/C ratio above 1.5 is indicated by orange dots. **e)** Histogram of time between initial increase (N/C ratio >1.5) and peak N/C ratio in the presence and absence of PP2A-B55 (n>50).

To confirm this phosphatase-specific effect further, we deleted the major catalytic subunit of the PP2A holoenzyme (*ppa2*), which partially reduces PP2A-B55 activity. As expected, SynCut3-T19 is net phosphorylated marginally earlier in the absence of the major PP2A catalytic subunit (*ppa2*) (Extended Data Fig. 7b). These data show that PP2A-B55 plays a role in restricting this substrate from being net phosphorylated earlier during the cell cycle, thereby contributing to the rapidness of the phosphorylation level change at the G2/M transition.

### PP2A-B55 and CDC14 are rate-limiting for mitotic onset in the absence of Tyrosine15

Given that all four phosphatases oppose G2/M CDK substrate sites, we investigated whether they play a rate-limiting role for the onset of mitosis. The phosphatases might act as ‘thresholders’, restricting the net phosphorylation of multiple CDK substrate sites during interphase, until CDK activity has reached a specific threshold. This could influence the timing of mitotic onset. In fission yeast, cell length at cell division can be used as a proxy for cell cycle timing (Fig. 6a). As previously reported, in the absence of PP2A-B55 and CDC14, cells are accelerated into mitosis and thus divide at a shorter length^44,45^ (Fig. 6b). Cell length at division is not reduced in the absence of PP2A-B56^44^, and in the absence of PP1, cells rapidly cease to grow in length, as PP1 activity is essential for polarised cell growth^46^. Thus, cell length cannot be used as a proxy of cell cycle stage in the absence of PP1.

**Figure 6:**
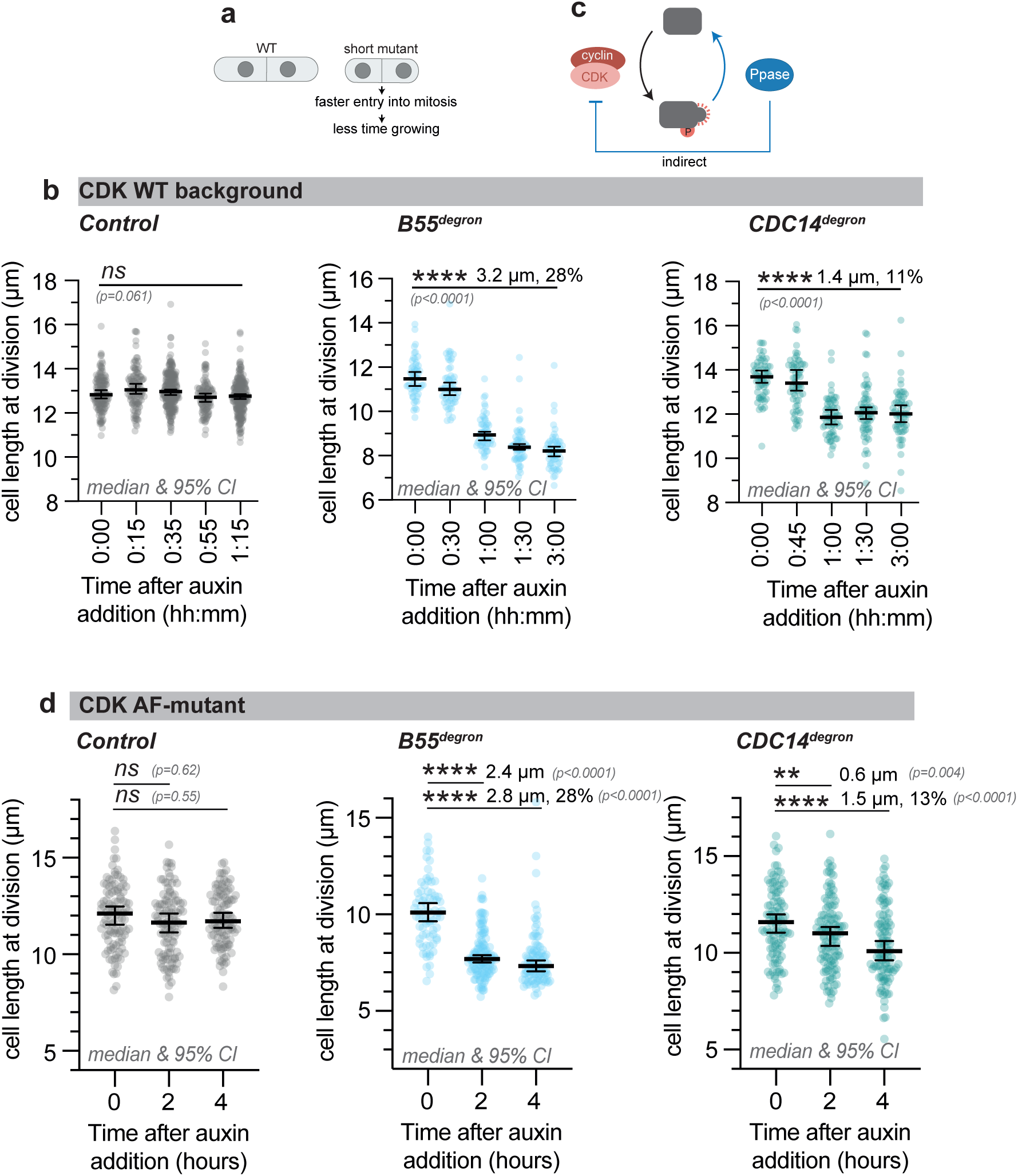
Degradation of PP2A-B55 and CDC14 accelerates cells into mitosis, independently of CDK activity regulation via the Tyrosine15 feedback loop. **a)** Schematic of cell length at division of fission yeast cells. **b)** Cell length at division of a *WT*, *TIR1 B55-sAID* and *TIR1 CDC14-sAID* strain at the indicated time after auxin addition. **c)** Schematic of the effect of phosphatase on CDK substrate phosphorylation either directly by dephosphorylating CDK substrate sites or by affecting CDK activity. **d)** Cell length at division of an *AFas CDKΔ CyclinBΔ TIR1 (n=109)*, *AFas CDKΔ CyclinBΔ TIR1 B55-sAID (n=57) and AFas CDKΔ CyclinBΔ TIR1 CDC14-sAID (n=62)* strain at the indicated time after auxin addition. **b,d)** Error bars represent median and 95% CI. Student’s t-test (two-tailed, unpaired) was used to determine statistical significance.

PP2A-B55 and CDC14 are rate-limiting for mitotic onset, which could either be because the phosphatases dephosphorylate CDK substrates or they regulate CDK activity via the Tyrosine15 feedback loop^47,48^ (Fig. 6c). To distinguish these two possibilities, we tested whether an advancement into mitosis is also seen in a CDK mutant background that is not subject to the Tyrosine15 feedback loop (CDK^AF^ mutant)^49^. In the CDK^AF^-mutant background, degradation of PP2A-B55 resulted in a 28 % reduction in cell length at division, while degradation of CDC14 led to a 13 % reduction (Fig. 6d). This indicates that both phosphatases influence the timing of the G2/M transition in a manner that is independent of the regulation of CDK activity through the Tyrosine15 feedback loop. As a control, we also repeated this experiment with the tyrosine phosphatase Pyp1, which negatively affects CDK activity, by regulating the Tyrosine15 feedback loop via the Sty1 kinase^50,51^. Cells lacking Pyp1 are therefore accelerated into mitosis^44^. Consistent with this, degradation of Pyp1 in a wild-type CDK background reduced cell length at division (Extended Data Fig. 8a). However, degradation of Pyp1 in a CDK^AF^-mutant background did not lead to a reduction in cell size (Extended Data Fig. 8b). This indicates that Pyp1’s negative effect on the G2/M transition does not function independently of the Tyrosine15 feedback loop, unlike PP2A-B55 and CDC14.

We degraded PP2A-B55 and CDC14 in an asynchronous culture predominantly containing cells in G2 phase and mitosis. The time between the degradation of the phosphatase and the reduction in cell length indicates when during the cell cycle the phosphatase exerts its negative regulatory effect. The later after the degradation of the phosphatase the reduction in length starts, the earlier this effect occurs during the cell cycle. Two hours after PP2A-B55 degradation, cell length at division was reduced by 24 %, while two hours after CDC14 degradation, cell length at division was only reduced by 5 % (Fig. 6d). This suggests that the rate-limiting effect of CDC14 occurs at an earlier stage of the cell cycle than that of PP2A-B55, consistent with the earlier result that CDC14 substrates are, on average, net phosphorylated earlier during the cell cycle (Extended Data Fig. 8c).

Our data show that PP2A-B55 and, to a lesser extent, CDC14, influence the timing of the G2/M transition independently of CDK activity regulation via Tyrosine15. This suggests that in a CDK^AF^ mutant background in the absence of PP2A-B55 and CDC14, critical CDK/PP2A-B55 and CDK/CDC14 substrates are phosphorylated earlier in G2, thereby advancing cells into mitosis.

## Discussion

Our dataset provides a comprehensive resource of CDK-dependent phosphatase substrates important for eukaryotic cell cycle regulation *in vivo*. We identified substrates of four different phosphatases in fission yeast (PP2A-B55, PP2A-B56, PP1, and CDC14), which together target 59% of identified CDK substrate sites (Fig. 2b). Of these sites, over 90 % were targeted by just one of the investigated phosphatases (Extended Data Fig. 4g), indicating that there is a division of labour between the phosphatases. Only around 5 % of CDK substrate sites were targeted non-redundantly by two or three different phosphatases, such that the degradation of each of the phosphatases reduced the observed dephosphorylation rate of a specific site upon CDK inhibition. We did not identify an opposing phosphatase for 41 % of CDK substrate sites. This may be because other phosphatases, such as PP4 or PP6^7,52,53^, dephosphorylate these sites. Alternatively, these sites may be targeted redundantly by multiple phosphatases, such that the degradation of one phosphatase is fully compensated by another phosphatase, and so no difference in dephosphorylation rate is observed upon depletion of any of the investigated phosphatases. Furthermore, we identified a total of around 3,600 CDK-independent phosphatase substrates, which highlights the fact that a single phosphatase can dephosphorylate diverse kinase substrates, thereby affecting diverse cellular processes (Supplementary table 2).

We have shown that CDK substrate sites dephosphorylated by CDC14 and PP2A-B56 are, on average, net phosphorylated earlier in G2, followed in order by CDK substrate sites dephosphorylated by PP1 and PP2A-B55. Thus, the identity of the phosphatase that targets a CDK substrate site influences the phosphorylation timing of the CDK substrate during progression through G2 into mitosis. Our data, therefore, indicate that the division of labour between the phosphatases in dephosphorylating different CDK substrates plays a role in regulating the net phosphorylation timing of CDK substrates. Increasing the turnover rate of a phosphosite can delay its net phosphorylation to a later point in the cell cycle. Depleting the investigated phosphatases slowed the dephosphorylation of CDK substrate sites after CDK inhibition, on average, by 3 times and, for certain substrates, up to 10 times (Extended Data Fig. 3c), suggesting that phosphatase activity has the capacity to affect the timing of net substrate phosphorylation via phosphosite turnover rates. The identity of the phosphatase is one of several determinants that influence the timing and phosphorylation pattern of CDK substrate sites, including CDK substrate sensitivity to CDK activity, SLiMs, multisite phosphorylation, and the subcellular localisation of the substrate^15,40–42^. A combination of these factors will determine the precise timing of each CDK substrate phosphorylation, and the weight of each factor is likely to vary between individual CDK substrate sites (Fig. 7a).

**Figure 7:**
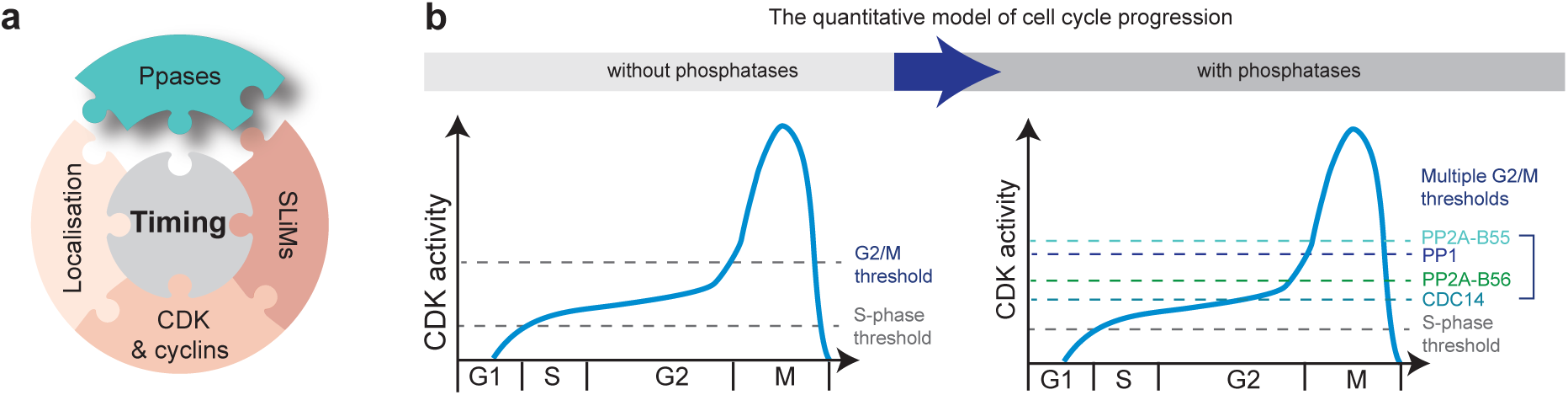
The role of CDK-opposing phosphatases in regulating the timing of CDK substrate phosphorylation. **a)** Schematic of the different factors influencing the phosphorylation timing of CDK substrate sites**. b)** Schematic of the effect of phosphatases on the threshold of the quantitative model of cell cycle progression.

The difference in phosphorylation timing between the groups of phosphatase substrates may be due to differing catalytic efficiencies of the phosphatases. Alternatively, this difference in time may be achieved by the activity of phosphatases being downregulated at different points during the cell cycle. CDK-opposing phosphatases are active during interphase and their activities are downregulated to different extents during mitosis^7^. At mitotic exit, PP1 is reactivated first and then reactivates PP2A-B55^54,55^. A similar mechanism could potentially operate in reverse at the G2/M transition. Different phosphatases becoming downregulated at different points during the G2/M transition would allow their respective CDK substrate sites to be net phosphorylated at different times in the cell cycle.

The quantitative model of the cell cycle proposes that progression through the cell cycle is ordered by an increase in net CDK activity, with cell cycle events happening at different net CDK activity thresholds^2,56^ (Fig. 7b). The fact that CDK substrates, which are dephosphorylated by different phosphatases, are net phosphorylated at different times (Fig. 4a), would allow finer tuning of phosphorylation times during G2 and mitosis. This is important given the complexity of the transition from G2 into mitosis when many CDK substrates need to be phosphorylated in a precise order while CDK activity is rising rapidly^2,57,58^. Our data imply that rather than there being one mitotic threshold, there are multiple phosphorylation thresholds at the G2/M transition for different groups of CDK substrates targeted by different phosphatases (Fig. 7b). The division of labour between phosphatases may, therefore, provide a finer time resolution at the G2/M transition, allowing for a more precisely ordered phosphorylation of hundreds of G2/M CDK substrates.

It remains unclear which of these CDK substrates are essential for entry into mitosis. This is highlighted by our result that in the absence of PP2A-B55 and, to a lesser extent, CDC14, cells are accelerated into mitosis independently of CDK activity regulation via the Tyrosine15 feedback loop (Fig. 6d). This advancement is likely due to earlier net phosphorylation of a particular set of critical CDK substrates and could either result from PP2A-55 and CDC14 targeting different sites on the same critical proteins, or PP2A-B55 and CDC14 targeting different substrate proteins, which can independently advance cells into mitosis. We identified 21 proteins which contain both PP2A-B55 substrate sites and CDC14 substrate sites and might, therefore, be rate-limiting for mitotic entry. They include proteins involved in chromosome segregation (INCENP^Pic1^, Survivin^Bir1^), chromatin structure (Spt2, Cph1), transcription regulation (Bdp1) and nuclear-cytoplasmic transport (Nup60). Both phosphatases target other potentially rate-limiting substrates, which may independently advance cells into mitosis, such as the Condensin subunit Cut3, which is targeted by PP2A-B55 (Supplementary Table 1). We propose that both these phosphatases act as thresholders, restricting the phosphorylation of CDK substrate sites which are critical for mitotic entry. Consistent with our data, deletions of CDC14 and the major catalytic subunit of PP2A have been shown to make cells more resistant to a decrease in global CDK activity^59^, likely because in the absence of these phosphatases, CDK can phosphorylate critical substrates earlier. Similarly, in budding yeast, PP2A-B55 acts as a thresholder, delaying the phosphorylation of threonine sites during the cell cycle^16^. With our experimental setup, we could not investigate the effect of PP1 on mitotic entry. However, removing PP1 from the centrosome allows an S-phase cyclin-CDK complex, which can normally only drive S-phase, to drive mitosis in fission yeast^3^. This suggests that deleting PP1 sufficiently increases the activity of the S-phase CDK to drive mitosis.

We conclude that the temporal order of cell cycle events brought about by CDK phosphorylating hundreds of substrates cannot be understood solely by considering the protein kinase but must also take account of the counteracting activities of at least the four investigated phosphatases: PP2A-B55, PP2A-B56, PP1, and CDC14. Our conclusions are likely relevant for core cell cycle control in other eukaryotes, given the high degree of conversation of core cell cycle architecture and substrate specificity of the phosphatases.

## Acknowledgements

We thank J. Curran, J. Greenwood, B. Whyte, S. Willich, T. Hammond and N. Kapadia for their comments on the manuscript, J. Curran for help with plasmid construction, T. Carr and A. Watson for *S. pombe* strains with the OsTIR1(F74A)-NLS construct and L. Du for the auxin-tagging plasmid. This work was supported by the Francis Crick Institute, which receives its core funding from Cancer Research UK (CC2003), the UK Medical Research Council (CC2003), and the Wellcome Trust (CC2003). In addition, this work was supported by the Wellcome Trust Grant to P.N. (grant number 214183), The Lord Leonard and Lady Estelle Wolfson Foundation, and Woosnam Foundation. T.U.Z. received funding from the Boehringer Ingelheim Fonds. For the purpose of Open Access, the author has applied a CC BY public copyright licence to any Author Accepted Manuscript version arising from this submission.

## Contributions

T.U.Z. and P.N. initiated the study. T.U.Z. designed and performed all experiments. T.A. and T.U.Z. performed mass spectrometry experiments. T.U.Z. performed data analysis. T.U.Z. and P.N. wrote the manuscript with input from all authors.

## Methods

### Schizosaccharomyces pombe genetics and cell culture

Fission yeast media and growth conditions have been described previously^60^. Strains were constructed by transformation or mating as previously described^60^ and checked for correct genotype by PCR (Supplementary Table 3). All experiments were performed with exponentially growing cultures in YE4S media (yeast extract with the supplements adenine, leucine, histidine, and uridine at 0.15 g/L) at 32 °C unless otherwise stated. Experiments with strains including the *cdc2(AFas)-cdc13 TIR1* background were performed in YE4S supplemented with 3 % w/v galactose to prevent flocculation^61^.

### Fluorescence microscopy

All microscopy experiments were performed using a Nikon Ti12 inverted microscope with Perfect Focus System, an Okolab environmental chamber and a Prime BSII sCMOS camera (Photometrics). Cells were imaged using an Olympus 100X objective lens with immersion oil (NA 1.45). The microscope was controlled with Micro-Manager v2.0 software (Open-imaging)^62^. For time-lapse imaging, cells were plated onto thin agarose pads, adapted from previous work in S. *cerevisiae*^63^. Cell masks were generated in Matlab and timelapse experiments analysed as previously described^64^.

### Protein extraction and western blotting

Protein extracts were prepared by adding ice-cold trichloroacetic acid (100 %, w/v) to cell culture samples to a final volume of 10 %. Cells were kept on ice for at least 30 min, pelleted at 3,000 g, and washed in ice-cold acetone, before dry pellets were stored at -80 °C. Pellets were washed and resuspended in lysis buffer (8 M urea, 50 mM ammonium bicarbonate, 5 mM EDTA (Ethylenediaminetetraacetic acid), 1 mM phenylmethylsulfonyl fluoride, protease inhibitor cocktail set III, and PhosSTOP phosphatase inhibitor cocktail). Acid wash glass beads (0.4 mm) were added to cell suspensions and beaten (using FastPrep120) to break cells. Cell debris was pelleted out and supernatant recovered as a protein extract (stored at -80 °C).

Protein detection by western blotting was performed for AID using a 1:1000 dilution of anti-mAID (mouse monoclonal antibody, MBL, M214-3), blocked with 5 % milk in TBS-T, and for TPxK-phopshorylation using a 1:1000 dilution of anti-pTPxK antibody (rabbit monoclonal, Cell Signalling Technology, D9V5N), blocked in 5 % BSA in TBS-T. The secondary antibodies used in this study were: 1:5000 mouse IgG HRP-linked whole antibody from sheep (GE Healthcare Cat# NA931, RRID:AB_772210); 1:5000 rabbit IgG HRP-linked whole antibody from donkey (GE healthcare, Cat# NA934, RRID:AB_772206). Signal was detected using SuperSignal West Femto Maximum Sensitivity Substrate (34095, Life Technologies) and imaged on an Amersham Imager 600.

### Tandem mass tag proteomics

Each protein sample (300 µg) was reduced with 5 mM dithiothreitol (DTT) for 25 min at 56 °C, alkylated with 10 mM iodoacetamide (30 min, room temperature, dark) and then quenched with 7.5 M DTT. Samples were digested using S-Traps (Protifi) spin columns, with the variation that 50 mM HEPES (pH 8.5) was used in place of ammonium bicarbonate. The digested samples were labelled with the TMTpro 16plex Isobaric Label Reagent Set (Thermo Fisher) according to the manufacturer’s instructions. After labelling and mixing, multiplexed samples were desalted using a Thermo desalting column. Phosphopeptide enrichment was performed using sequential enrichment from metal oxide affinity chromatography (SMOAC, Thermo Fisher). Initial enrichment was done using the HighSelect TiO2 Phosphopeptide Enrichment kit followed by the HighSelect Fe-NTA Phosphopeptide Enrichment kit (both from Thermo Scientific). Phosphopeptides and non-bound flow-through fractions were desalted and fractionated using the High-pH Reversed-Phase Peptide Fractionation kit (Pierce) and analysed on Orbitrap Eclipse Tribrid mass spectrometer (Thermo Fisher) coupled to an UltiMate 3000 HPLC system for online liquid chromatography separation. Each run consisted of a 3-hour gradient elution from a 75 μm × 50 cm C18 column operated in Data-dependent acquisition mode.

### Mass spectrometry data analysis

MaxQuant (version 1.6.14.0 and 2.4.7.0) was used to search data against an *S. pombe* proteome FASTA file, extracted from PomBase and amended to include common contaminants. Default MaxQuant parameters were used apart from adding phospho(STY) as a variable modification for phosphopeptide-enriched samples. Phospho(STY).txt (phosphopeptides) and ProteinGroup.txt (peptides) MaxQuant outputs were imported into Perseus (version 1.6.14.0) for further analysis. Data was filtered for valid values (100 %), localisation probability (>0.75) and median normalised. The same phosphosite with a different phosphorylation multiplicity was treated as a separate phosphorylation event. All further analyses, including curve fitting and plotting of heatmaps, were done using Matlab (R2024b) and RStudio. Phosphatase substrates (proteins and sites) are listed in Supplementary Table 1 (CDK-dependent phosphatase substrates) and 2 (CDK-independent phosphatase substrates). Enrichment logos were made using iceLogo^65^, GO enrichment analysis was conducted using ShinyGO^66^ and the disordered regions of the *S. pombe* proteome were searched for SLiMs using SLiMSearch4^67^. The phosphoproteomic timecourse analysed in Fig. 4 and Extended Data Fig. 6 was previously described^2^. In brief, a Cdc13-Cdc2-as strain in which Cig1, Cig2, Puc1 had been deleted (ΔCCP), was grown in EMM4S + SILAC (stable isotope labelling with amino acids in cell culture) media at 32 °C. Cells were arrested at G2 using 1-NmPP1 and released into mitosis by washing into 1-NmPP1-free media^2^.

### Phosphatase substrate identification

The phosphorylation of phosphosites, normalised to timepoint 0 (mean of two technical replicates was used for t0), was used to identify CDK substrate sites and CDK-dependent phosphatase substrates. Sites that decreased by 50% in their phosphorylation level within 12 minutes after t0 were included. An exponential decay function was fitted to the data using the non-linear least-squares method in Matlab, calculating the dephosphorylation rate constant and half-life. The criteria used to define CDK substrate sites and CDK-dependent phosphatase substrates are listed in Extended Data Fig. 1c. For CDK-independent phosphatase substrates, a linear regression was fitted to the data (not normalised) of the control and -Ppase condition using RStudio. Sites were defined as CDK-independent if the slope was >-0.05 and < 0.05 and the RMSE <0.4. A student’s t-test (corrected for multiple testing using the Benjamini-Hochberg method) was used to determine statistical difference between the mean phosphorylation level of the Control and -Ppase condition (see Extended Data Figure 5a).

## Data representation

All statistical tests were conducted using RStudio 2024b or GraphPad Prism 10. The central point for data given is either the mean value, with whiskers indicating 95 % confidence intervals, or the median value, with whiskers indicating IQR, unless otherwise stated.

## Data and code availability

All mass spectrometry data generated have been deposited to the ProteomeXchange Consortium via the PRIDE partner repository with dataset identifier XXXX. Mass spectrometry sample names are listed in Supplementary table 4. Previously described custom scripts used to analyse fluorescent time-lapse data can be found at https://github.com/nkapadia27/Spatiotemporal-Orchestration-of-Mitosis^64^.

## Extended Data Figures

**Extended Data Figure 1:**
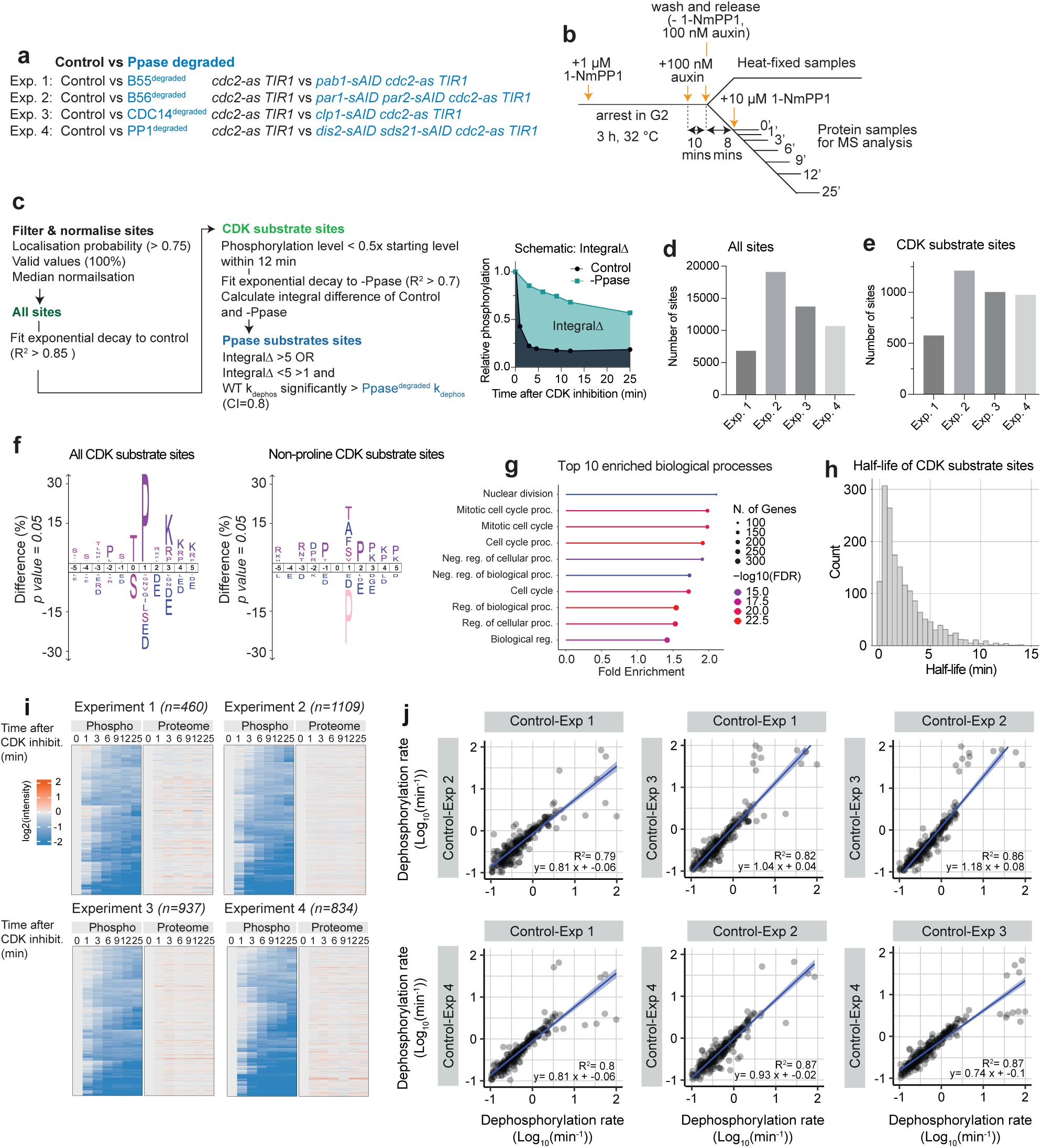
Identification of *in vivo* CDK substrate sites. **a)** List of strains used in each of the four phosphoproteomic experiments. **b)** Schematic of phosphoproteomic experiment. **c)** Left: Criteria for classifying phosphatase substrates. Right: Schematic plot showing the difference of integrals (from 0-25 mins after CDK inhibition), used to classify phosphatase substrates. **d-e)** Bar graph representing the number of d) all phosphosites detected in the four phosphoproteomic experiments after filtering and e) number of identified CDK substrate sites. **f)** IceLogo representation of over- and underrepresented amino acid residues surrounding the phosphorylation sites of all identified CDK substrate sites (left) and all CDK substrate sites which do not contain a Proline at the +1 position (right, non-consensus sites)^65^. **g)** GO-enrichment of CDK substrate sites, showing the 10 most enriched biological processes using ShinyGO^66^. **h)** Histogram of half-life (min) of all identified CDK substrate sites. **i)** Heatmap visualising the changes in phosphosites (left) and the respective proteins (right) upon inactivation of CDK activity. Only sites which are classified as CDK substrate sites and for which the corresponding protein was detected in the proteome are plotted. **j)** Pairwise linear regressions of dephosphorylation rates determined for phosphosites in the control conditions of the four different experiments.

**Extended Data Figure 2:**
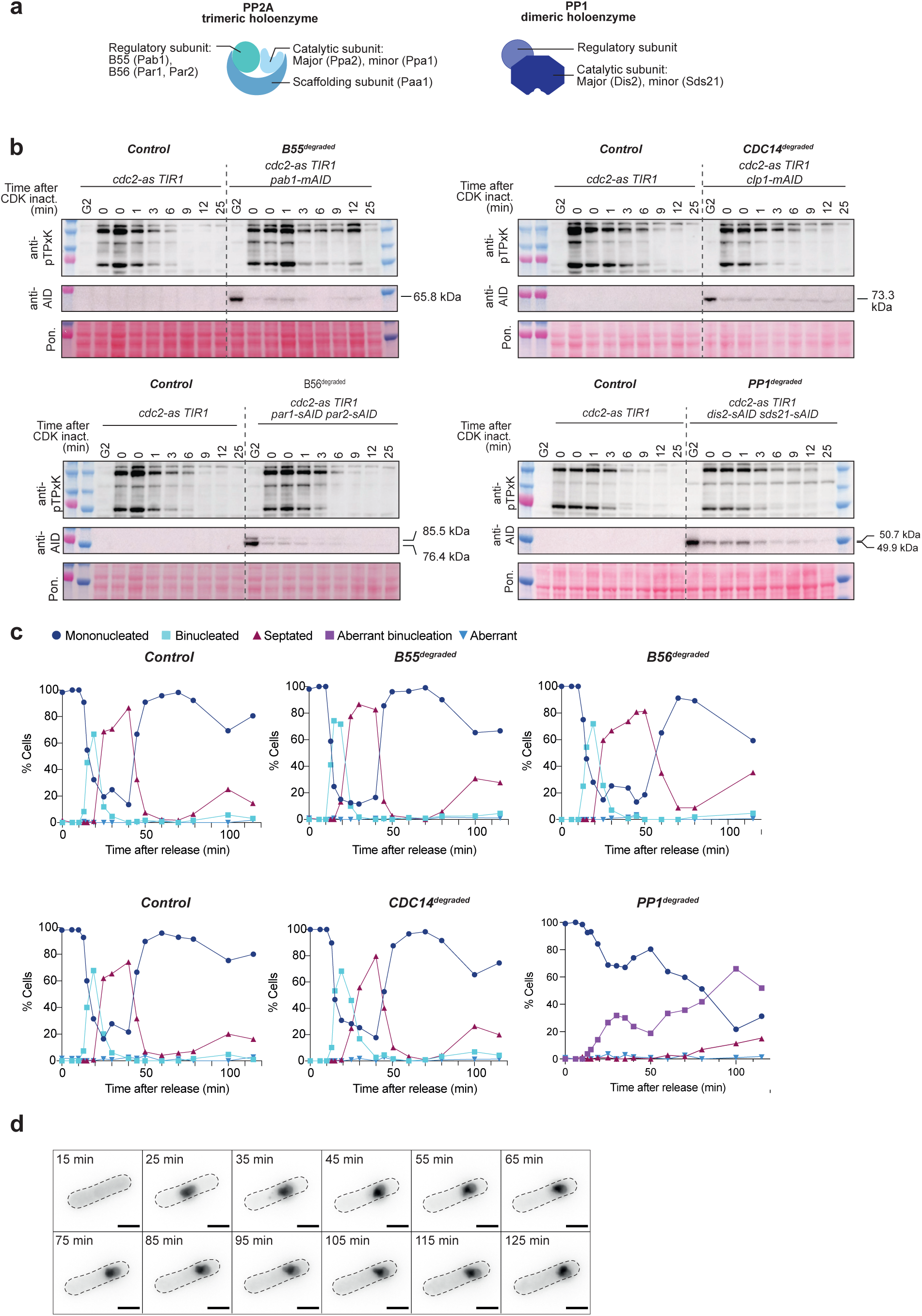
Degradation of phosphatases using the AID system. **a)** Schematic of PP2A and PP1 holoenzyme subunits. **b)** Western blots of phospho-CDK substrates and sAID-tagged phosphatases after release from G2 arrest. Numbers indicate the time in minutes after CDK inactivation at peak mitosis. Two samples were taken simultaneously at timepoint 0. Membranes were probed with an anti-pTPxK antibody and anti-AID antibody. Ponceau-S stain was used for total protein normalisation. **c)** Binucleation and septation indices were quantified from DAPI- and calcofluor-stained fixed cells. Timepoint zero represents the point at which G2 arrested cells were washed and released into 1-NmPP1-free media. At least 100 cells were counted per timepoint. Degradation of PP2A-B56 regulatory subunits Par1 and Par2 led to broader septation peaks and a delay in cytokinesis, consistent with Par1 promoting cytokinesis after a prolonged metaphase arrest^68^ and *par1Δpar2Δ* double mutants showing a higher incidence of double septa and misplaced septa^69^. Degradation of CDC14 resulted in a short delay in the start of septation, consistent with CDC14 playing a role in cytokinesis via the septation initiation network^70,71^. Degradation of both catalytic subunits of PP1 (Dis2 and Sds21) led to a severe binucleation defect (aberrant binucleation; see also Extended Data Fig. 2d), consistent with PP1 activity being required for accurate chromosome segregation^54^. **d)** Example timelapse images of SynCut3-mCherry fluorescence (Cut3 is a Condensin subunit and is used here as a nuclear marker) in *TIR1 dis2-sAID sds21-sAID cdc2-as syncut3-mCherry* cells. Cells lacking PP1 were arrested in G2 using 1 µm 1-NmPP1 and released into mitosis, revealing an asymmetric division of the nucleus. Time in images refers to minutes after release from G2. Cells are not viable long-term (data not shown), consistent with a *dis2Δsds21Δ* mutant being lethal^72^. The dashed lines indicate single cell masks, generated from the bright-field image. Scale bar represents 5 µm.

**Extended Data Figure 3:**
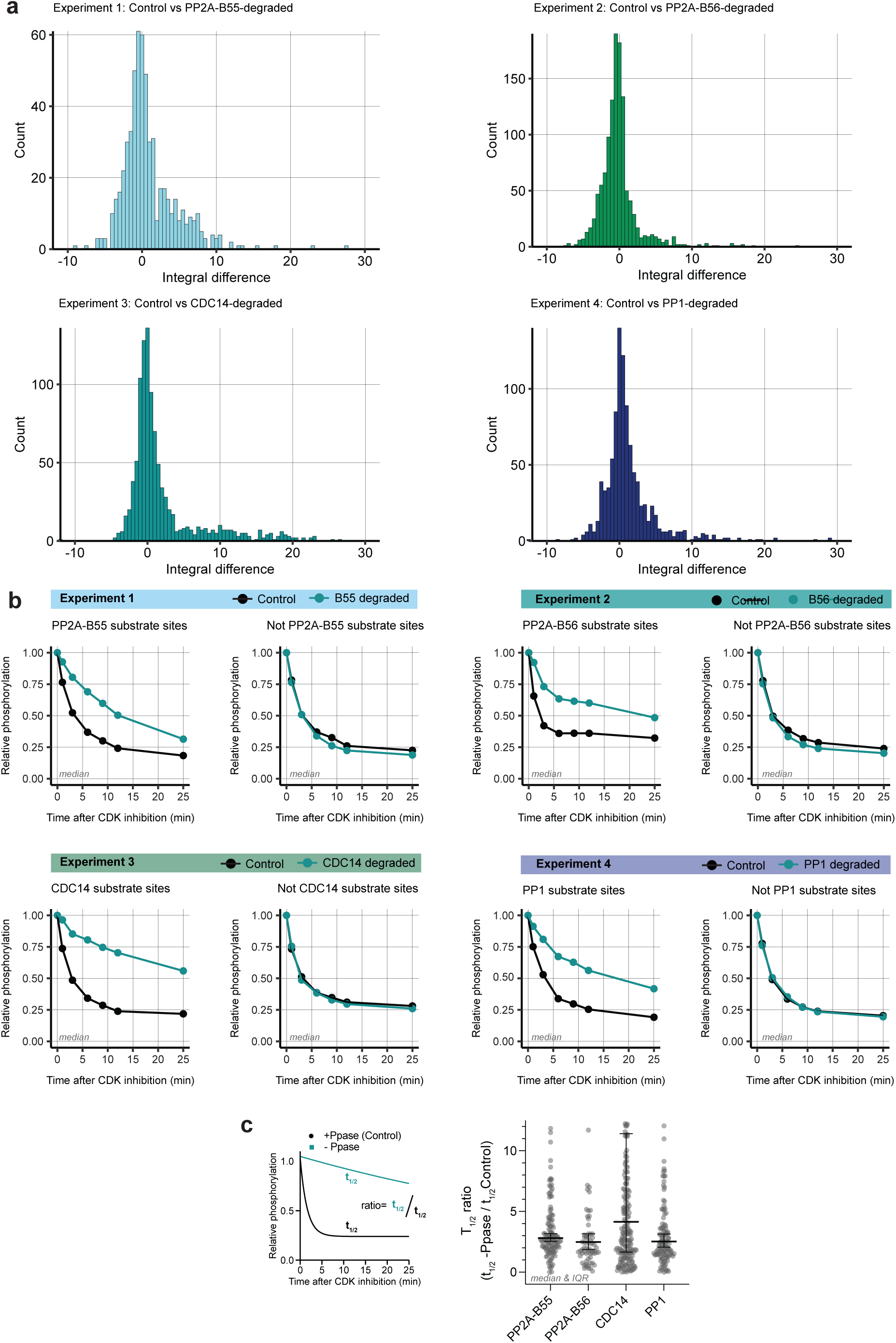
Identification of *in vivo* phosphatase substrates. **a)** Histogram of integral difference (explained in Extended Data Fig. 1c) between -Ppase and Control condition in all CDK substrate sites in the four different phosphoproteomic experiments. **b)** Median phosphorylation of all phosphatase substrates and not phosphatase substrates in the presence and absence of the respective phosphatase. **c)** Left panel: Schematic of ratio of t_1/2_ of -Ppase and control condition. Right panel: Beeswarm plots portraying the ratio of the t_1/2_ of -Ppase and Control condition for PP2A-B55, PP2A-B56, CDC14, and PP1 substrates. Error bars denote the median and IQR.

**Extended Data Figure 4:**
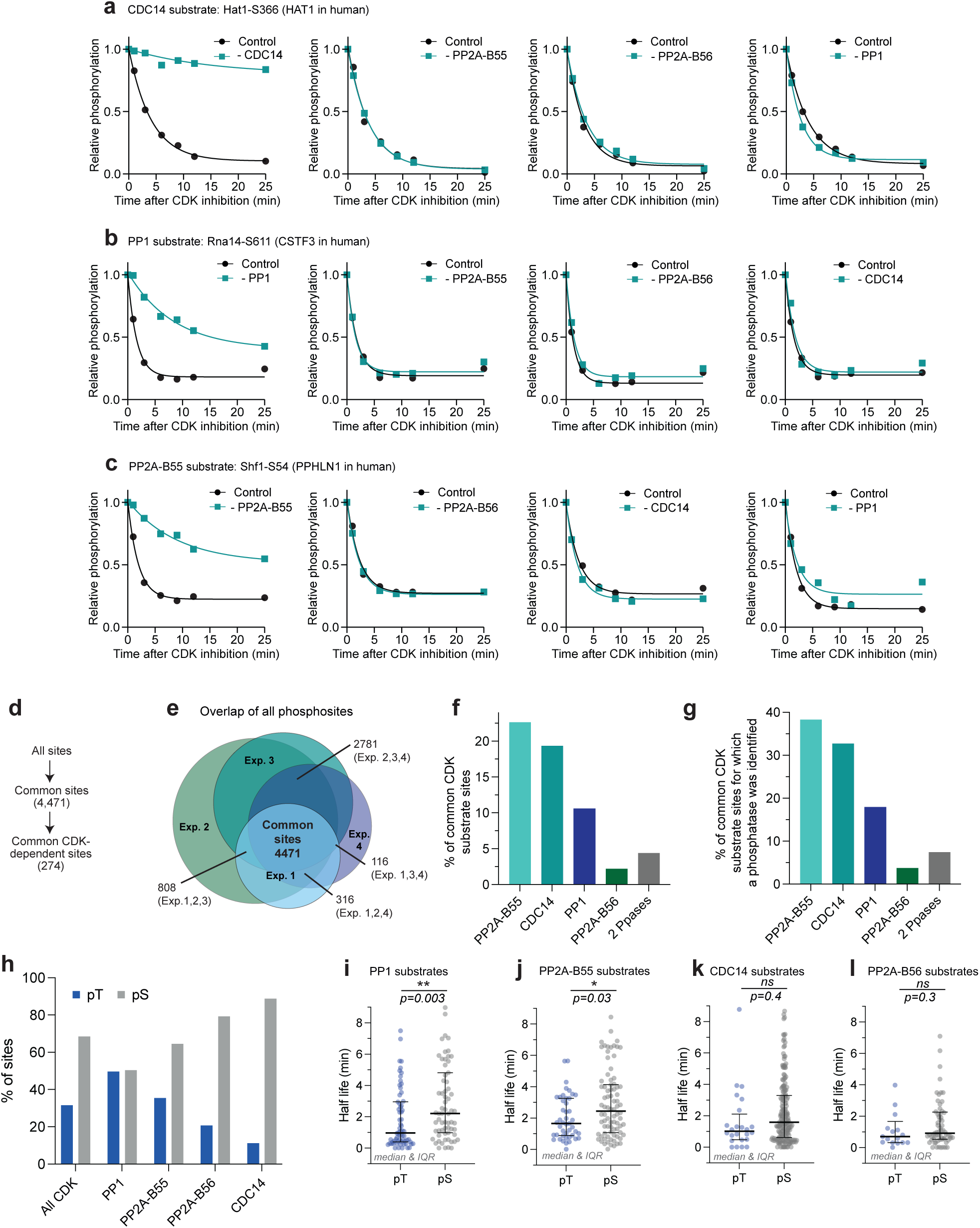
Substrate specificity of investigated phosphatases. **a-c)** Example of phosphatase substrates, which are targeted by just one of the identified phosphatase substrates. Relative phosphorylation level in the presence (black) and absence (turquoise) of phosphatase of interest upon CDK inhibition. Curves are a one-phase exponential decay fitted to the relative phosphorylation level determined by phosphoproteomics. **d)** Schematic of filtering for sites, which are identified in all 4 phosphoproteomic experiments.**e)** Venn diagram depicting the overlap of all detected phosphorylation sites in the four experiments. **f-g)** Bar graph showing the phosphatase substrates as a percentage of **f)** all common CDK substrate sites **g)** CDK-substrate sites that are targeted by at least one of the investigated phosphatases. **h)** Grouped bar graph representing the percentage of pThreonine and pSerine in CDK substrate sites and CDK-dependent phosphatase substrates. **i-l)** Half-life of pThreonine and pSerine CDK-dependent phosphatase substrates, for each of the four investigated phosphatase substrates. Error bars denote the median and IQR. Statistical difference between groups was determined using a Mann-Whitney test (two-tailed).

**Extended Data Figure 5:**
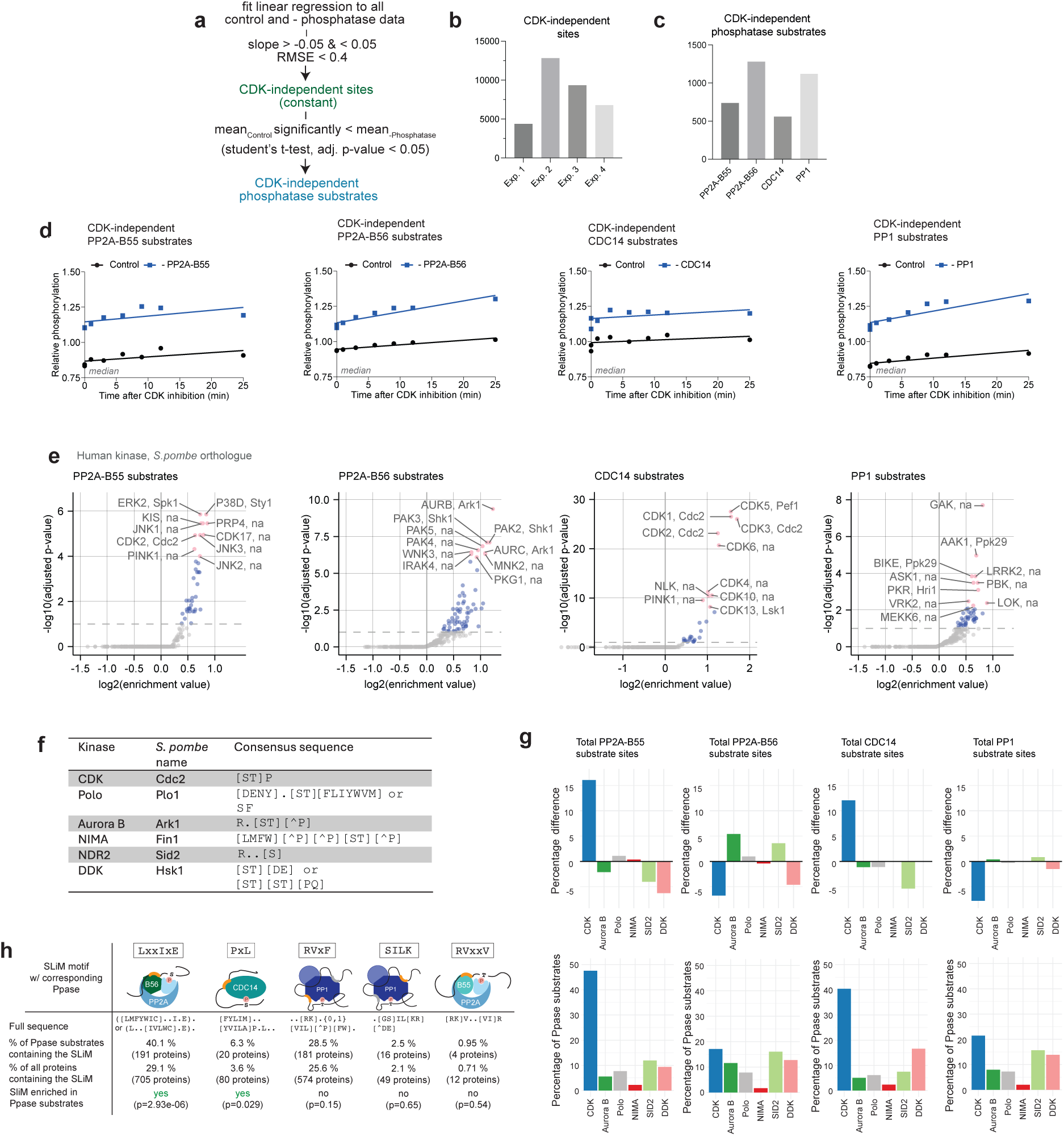
CDK-dependent and independent phosphatase substrates. **a)** Criteria to identify CDK-independent sites (constant sites) and CDK-independent phosphatase substrates. **b-c)** Bar graph representing the number of identified b) CDK-independent sites and c) CDK-independent phosphatase substrates in the four different phosphoproteomic experiments. **d)** Median phosphorylation of CDK-independent phosphatase substrates for PP2A-B55, PP2A-B56, PP1, and CDC14 in the presence (black) and absence (blue) of the respective phosphatases. A linear model was fitted through the median points. **f)** Motifs used to identify substrates of cell-cycle kinases. **g)** Motif enrichment analysis of phosphatase substrates for kinase motifs using the human kinome atlas^31^. Enrichments were determined using Fisher’s exact tests (corrected for multiple testing using the Benjamini-Hochberg method). **f)** Top panel: Bar graph representing the percentage difference of cell cycle kinase substrates present in the phosphatase substrates, compared to all phosphosites. Bottom panel: Top panel: Bar graph representing the percentage cell cycle kinase substrates present in the phosphatase substrates. **i)** Occurrence of SliMs in phosphatase substrates (CDK-dependent and independent)^38,39,73–75^. Enrichments were determined using Fisher’s exact tests.

**Extended Data Figure 6:**
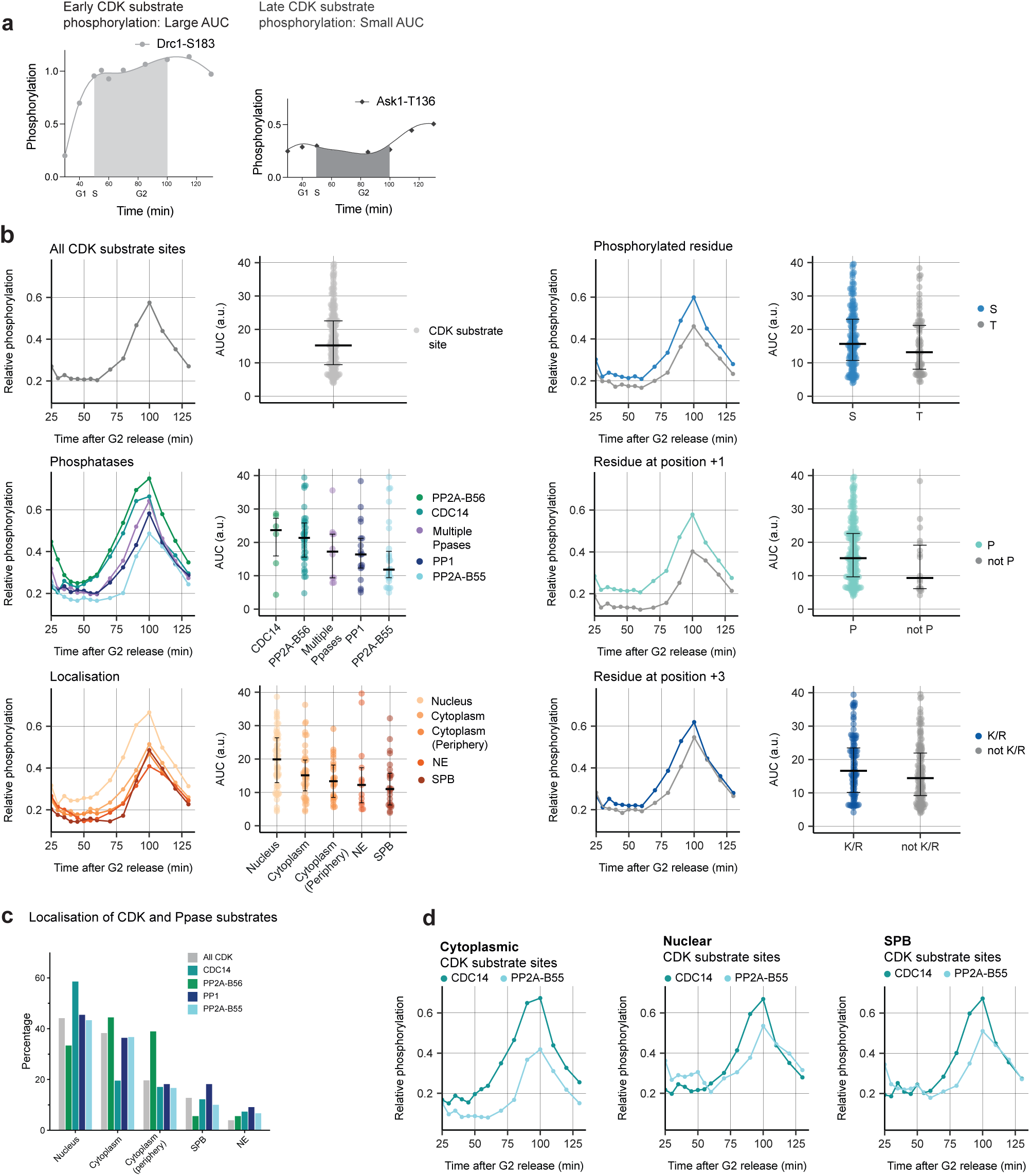
Estimating the effect of different determinants on the phosphorylation timing of CDK substrate sites. **a)** Example traces of relative phosphorylation of an example Early (Drc1-S183) and Late (Ask1-T136) CDK substrates throughout a cell cycle. Data are from a synchronised phosphoproteomics timecourse experiment^2^. AUC was calculated as a measure of phosphorylation timing within the cell cycle. **b)** Median relative phosphorylation (left) and AUC data (right) of CDK substrate sites during a synchronised cell cycle, split according to opposing phosphatase, localisation of the substrate, phosphorylated residue (S or T), Amino acid at position +3 and amino acid at position +1. Error bars in AUC plots denote the median and IQR. **c)** Grouped bar graph of sub-cellular localisations of phosphatase substrates. As a site can have multiple localisations, the percentages may not add up to 100. Fisher’s exact tests (corrected for multiple testing using the Benjamini-Hochberg method) were used to test whether phosphatase substrates were significantly enriched in any of the subcellular localisations and showed no significant difference for any of the groups (p>0.05). **d)** Median relative phosphorylation of CDC14 and PP2A-B55 CDK substrate sites during a synchronised cell cycle, filtered for sites which are localised in the cytoplasm (left panel), nucleus (middle panel) or SPB (right panel).

**Extended Data Figure 7:**
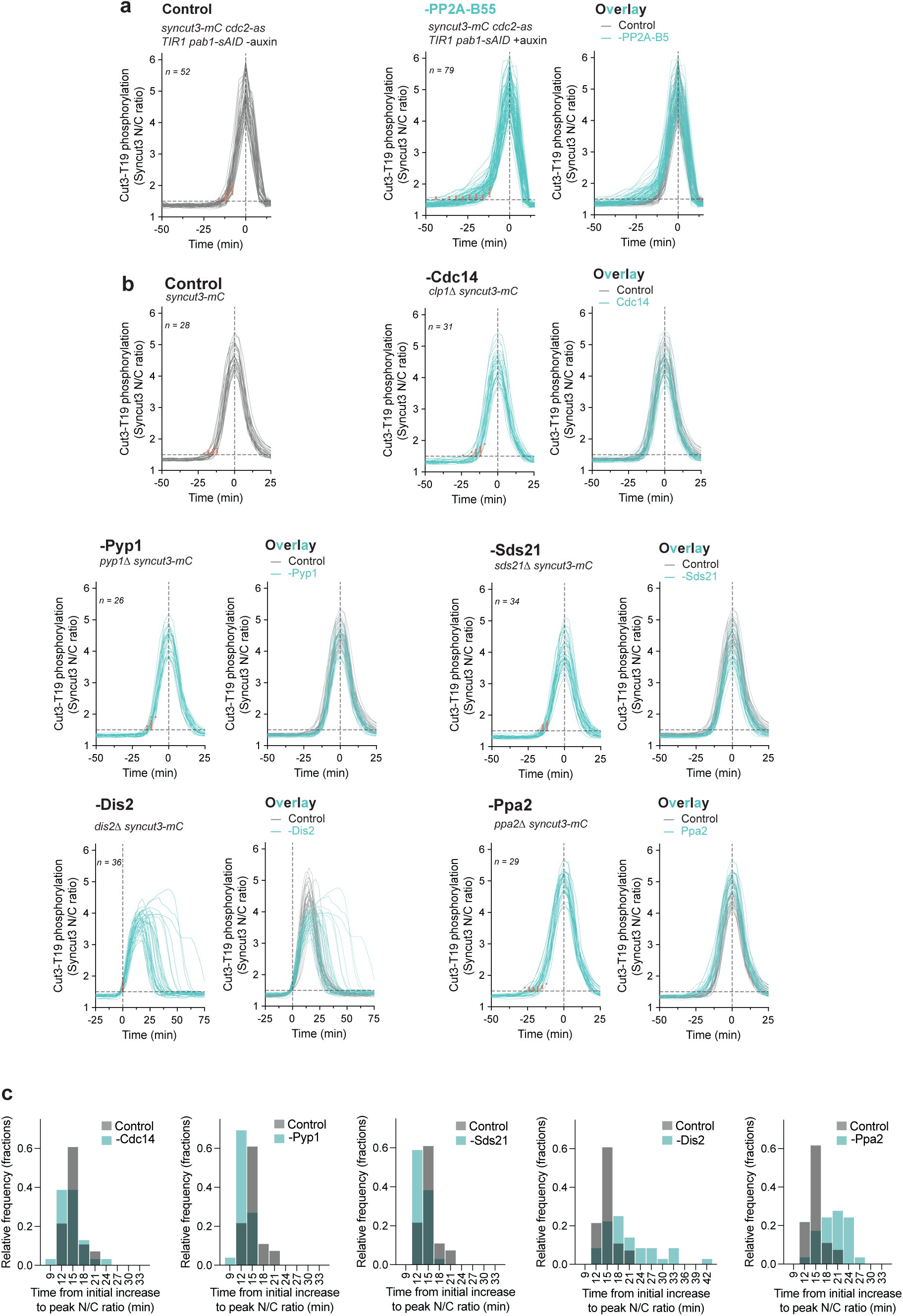
Effect of other cell cycle phosphatases on the phosphorylation pattern of SynCyt3. **a)** Single-cell traces of SynCut3-mC N/C ratio in the presence (control, grey) and absence of PP2A-B55 (B55 degraded, turquoise). Traces are aligned at peak N/C ratio (t=0) and the initial increase in N/C ratio above 1.5 is indicated by orange dots. Right panel shows an overlay of the control and - PP2A-B55 condition. **b)** Single-cell traces of SynCut3-mC N/C ratio in the presence (control, grey) and absence (Ppase-deleted, turquoise) of the indicated phosphatases. Traces are aligned at peak N/C ratio (t=0) and the initial increase in N/C ratio above 1.5 is indicated by orange dots. Right panel shows an overlay of the control and -PP2A-B55 condition. Cells lacking the major catalytic subunit of PP1 (Dis2) are aligned at initial increase (N/C>1.5). They show an aberrant mitotic progression, likely due to a delayed metaphase-anaphase transition (see also Extended Data Fig. 2c-d). **c)** Histograms of time between the initial increase of phosphorylation and peak N/C ratio in the presence (control, grey) and absence (Ppase-deleted, turquoise) of the indicated phosphatase. Experiments using the AID system shown in a were conducted at 32 °C, while experiments using Ppase-delete strains in were conducted at 25 °C. Thus the median time between initial increase and peak N/C ratio in the control takes 15 rather than 12 min (compare Fig. 5e and Extended Data Fig. 7c).

**Extended Data Figure 8:**
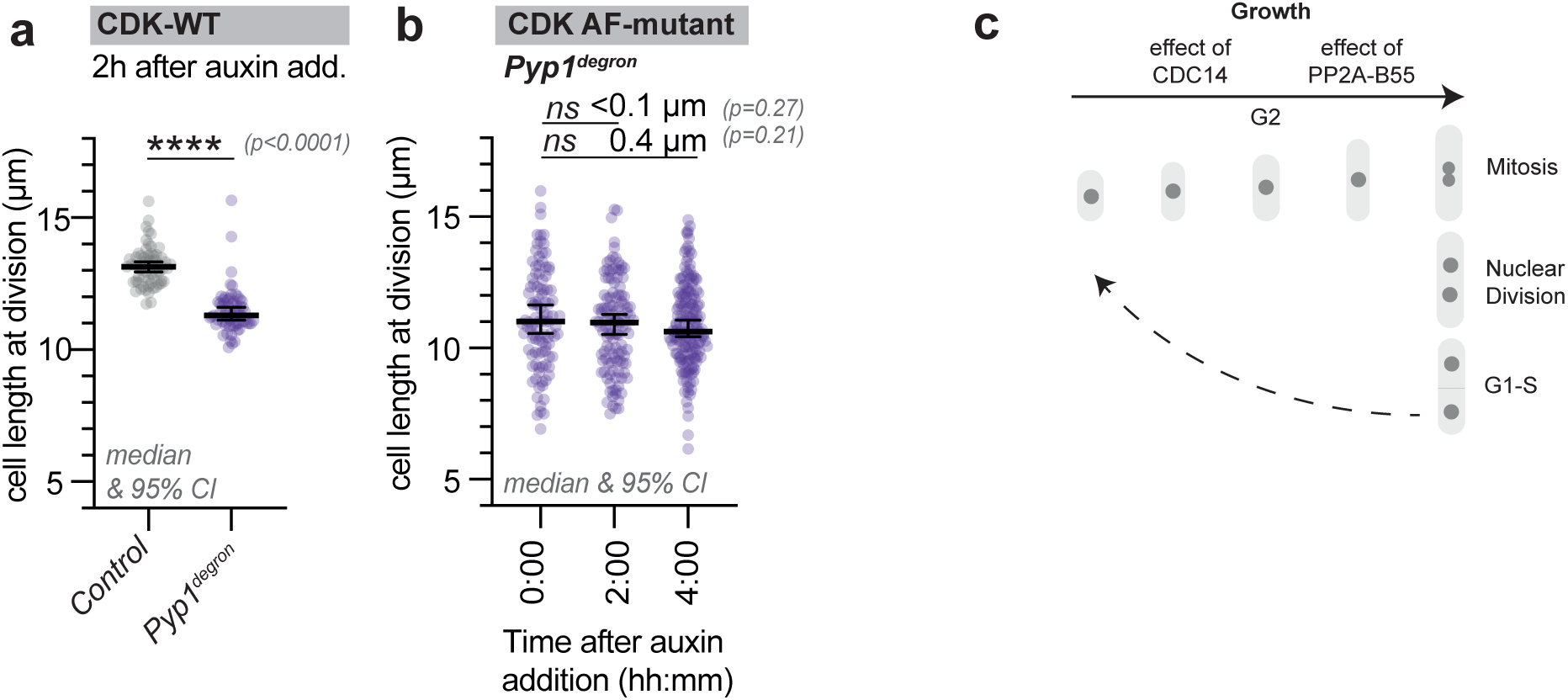
Pyp1’s delays mitotic onset by affecting CDK activity. **a)** Cell length at division of *WT*, and *pyp1Δ* cells (n=65). **b)** Cell length at division at the indicated time after auxin addition of an *AFas CDKΔ CyclinBΔ TIR1* and *AFas CDKΔ CyclinBΔ TIR1 pyp1-sAID* strain (n>100). a-b) Error bars represent median and 95% CI. Student’s t-test (two-tailed, unpaired) was used to determine statistical difference. **c)** Schematic of growth during an *S. pombe* cell cycle. The approximate times in G2 phase at which PP2A-B55 and CDC14 have their negative regulatory effect on the G2/M transition are indicated.

## References

1. Morgan, D. O. The Cell Cycle: Principles of Control. (New science press, 2007).

2. Swaffer, M. P., Jones, A. W., Flynn, H. R., Snijders, A. P. & Nurse, P. CDK Substrate Phosphorylation and Ordering the Cell Cycle. Cell 167, 1750–1761.e16 (2016).

3. Basu, S., Greenwood, J., Jones, A. W. & Nurse, P. Core control principles of the eukaryotic cell cycle. Nature 607, 381–386 (2022).

4. Gould, K. L. & Nurse, P. Tyrosine phosphorylation of the fission yeast cdc2+ protein kinase regulates entry into mitosis. Nature 342, 39–45 (1989).

5. Novak, B. & Tyson, J. J. Numerical analysis of a comprehensive model of M-phase control in Xenopus oocyte extracts and intact embryos. J Cell Sci 106, 1153–1168 (1993).

6. Loog, M. & Morgan, D. O. Cyclin specificity in the phosphorylation of cyclin-dependent kinase substrates. Nature 434, 104–108 (2005).

7. Nasa, I. & Kettenbach, A. N. Coordination of protein kinase and phosphoprotein phosphatase activities in mitosis. Front Cell Dev Biol 6, 1–14 (2018).

8. Nilsson, J. Protein phosphatases in the regulation of mitosis. Journal of Cell Biology 218, 395–409 (2019).

9. Castilho, P. V., Williams, B. C., Mochida, S., Zhao, Y. & Goldberg, M. L. The M phase kinase greatwall (Gwl) promotes inactivation of PP2A/B55δ, a phosphatase directed against CDK phosphosites. Mol Biol Cell 20, 4777–4789 (2009).

10. Mochida, S., Ikeo, S., Gannon, J. & Hunt, T. Regulated activity of PP2A-B55δ is crucial for controlling entry into and exit from mitosis in Xenopus egg extracts. EMBO J. 28, 2777–2785 (2009).

11. Wu, J. Q. et al. PP1-mediated dephosphorylation of phosphoproteins at mitotic exit is controlled by inhibitor-1 and PP1 phosphorylation. Nature Cell Biology 2009 11:5 11, 644–651 (2009).

12. Touati, S. A. et al. Cdc14 and PP2A Phosphatases Cooperate to Shape Phosphoproteome Dynamics during Mitotic Exit. Cell Rep 29, 2105–2119.e4 (2019).

13. Holder, J., Mohammed, S. & Barr, F. A. Ordered dephosphorylation initiated by the selective proteolysis of cyclin B drives mitotic exit. Elife 9, 1–33 (2020).

14. Holt, L. J. et al. Global analysis of Cdk1 substrate phosphorylation sites provides insights into evolution. Science (1979) 325, 1682–1686 (2009).

15. Uhlmann, F., Bouchoux, C. & López-Avilés, S. A quantitative model for cyclin-dependent kinase control of the cell cycle: revisited. Philosophical Transactions of the Royal Society B: Biological Sciences 366, 3572–3583 (2011).

16. Godfrey, M. et al. PP2ACdc55 Phosphatase Imposes Ordered Cell-Cycle Phosphorylation by Opposing Threonine Phosphorylation. Mol Cell 65, 393–402.e3 (2017).

17. Brautigan, D. L. Protein Ser/ Thr phosphatases - The ugly ducklings of cell signalling. FEBS Journal 280, 324–325 (2013).

18. Songyang, Z. et al. Use of an oriented peptide library to determine the optimal substrates of protein kinases. Curr Biol 4, 973–982 (1994).

19. Watson, A. T., Hassell-Hart, S., Spencer, J. & Carr, A. M. Rice (Oryza sativa) tir1 and 5′adamantyl-iaa significantly improve the auxin-inducible degron system in schizosaccharomyces pombe. Genes (Basel) 12, (2021).

20. Zhang, X. R. et al. An improved auxin-inducible degron system for fission yeast. G3: Genes, Genomes, Genetics 12, (2022).

21. Bollen, M., Peti, W., Ragusa, M. J. & Beullens, M. The extended PP1 toolkit: Designed to create specificity. Trends Biochem Sci 35, 450–458 (2010).

22. Cundell, M. J. et al. A PP2A-B55 recognition signal controls substrate dephosphorylation kinetics during mitotic exit. Journal of Cell Biology 214, 539–554 (2016).

23. Partscht, P., Simon, A., Chen, N. P., Erhardt, S. & Schiebel, E. The HIPK2/CDC14B-MeCP2 axis enhances the spindle assembly checkpoint block by promoting cyclin B translation. Sci Adv 9, (2023).

24. Sharma, K. et al. Ultradeep Human Phosphoproteome Reveals a Distinct Regulatory Nature of Tyr and Ser/Thr-Based Signaling. Cell Rep 8, 1583–1594 (2014).

25. Agostinis, P., Derua, R., Sarno, S., Goris, J. & Merlevede, W. Specificity of the polycation-stimulated (type-2A) and ATP, Mg-dependent (type-1) protein phosphatases toward substrates phosphorylated by P34cdc2 kinase. Eur J Biochem 205, 241–248 (1992).

26. Hoermann, B. et al. Dissecting the sequence determinants for dephosphorylation by the catalytic subunits of phosphatases PP1 and PP2A. Nat Commun 11, 1–20 (2020).

27. Kruse, T. et al. Mechanisms of site-specific dephosphorylation and kinase opposition imposed by PP2A regulatory subunits. EMBO J (2020) doi:10.15252/embj.2019103695.

28. Koch, A., Krug, K., Pengelley, S., Macek, B. & Hauf, S. Mitotic substrates of the kinase aurora with roles in chromatin regulation identified through quantitative phosphoproteomics of fission yeast. Sci Signal 4, 1–12 (2011).

29. Suzuki, K. et al. Identification of non-Ser/Thr-Pro consensus motifs for Cdk1 and their roles in mitotic regulation of C2H2 zinc finger proteins and Ect2. Sci Rep 5, 1–9 (2015).

30. Gray, C. H., Good, V. M., Tonks, N. K. & Barford, D. The structure of the cell cycle protein Cdc14 reveals a proline-directed protein phosphatase. EMBO J 22, 3524–3535 (2003).

31. Johnson, J. L. et al. An atlas of substrate specificities for the human serine/threonine kinome. Nature 613, 759–766 (2023).

32. Kettenbach, A. N. et al. Quantitative phosphoproteomics identifies substrates and functional modules of Aurora and Polo-like kinase activities in mitotic cells. Sci Signal 4, (2011).

33. Alexander, J. et al. Spatial exclusivity combined with positive and negative selection of phosphorylation motifs is the basis for context-dependent mitotic signaling. Sci Signal 4, 35– 37 (2011).

34. Lizcano, J. M. et al. Molecular basis for the substrate specificity of NIMA-related kinase-6 (NEK6). Evidence that NEK6 does not phosphorylate the hydrophobic motif of ribosomal S6 protein kinase and serum- and glucocorticoid-induced protein kinase in vivo. Journal of Biological Chemistry 277, 27839–27849 (2002).

35. Mah, A. S. et al. Substrate specificity analysis of protein kinase complex Dbf2-Mob1 by peptide library and proteome array screening. BMC Biochem 6, (2005).

36. Randell, J. C. W. et al. Mec1 Is One of Multiple Kinases that Prime the Mcm2-7 Helicase for Phosphorylation by Cdc7. Mol Cell 40, 353–363 (2010).

37. Nguyen, H. & Kettenbach, A. N. Substrate and phosphorylation site selection by phosphoprotein phosphatases. Trends Biochem Sci 1–13 (2023) doi:10.1016/j.tibs.2023.04.004.

38. Hertz, E. P. T. et al. A Conserved Motif Provides Binding Specificity to the PP2A-B56 Phosphatase. Mol Cell 63, 686–695 (2016).

39. Kataria, M. et al. A PxL motif promotes timely cell cycle substrate dephosphorylation by the Cdc14 phosphatase. Nat Struct Mol Biol 25, 1093–1102 (2018).

40. Basu, S. et al. The Hydrophobic Patch Directs Cyclin B to Centrosomes to Promote Global CDK Phosphorylation at Mitosis. Current Biology 30, 883–892.e4 (2020).

41. Örd, M. et al. Multisite phosphorylation code of CDK. Nat Struct Mol Biol 26, 649–658 (2019).

42. Kapadia, N. & Nurse, P. Spatiotemporal Orchestration of Mitosis by Cyclin-Dependent Kinase. Preprint at 10.1101/2024.04.29.591629 (2024).

43. Sutani, T. et al. Fission yeast condensin complex: Essential roles of non-SMC subunits for condensation and Cdc2 phosphorylation of Cut3/SMC4. Genes Dev 13, 2271–2283 (1999).

44. Hayles, J. et al. A genome-wide resource of cell cycle and cell shape genes of fission yeast. Open Biol 3, 130053 (2013).

45. Chica, N. et al. Nutritional control of cell size by the greatwall-endosulfine-PP2A·B55 pathway. Current Biology 26, 319–330 (2016).

46. Kokkoris, K., Castro, D. G. & Martin, S. G. The Tea4-PP1 landmark promotes local growth by dual Cdc42 GEF recruitment and GAP exclusion. J Cell Sci 127, 2005–2016 (2014).

47. Pomerening, J. R., Sun, Y. K. & Ferrell, J. E. Systems-level dissection of the cell-cycle oscillator: Bypassing positive feedback produces damped oscillations. Cell 122, 565–578 (2005).

48. Solomon, M. J., Glotzer, M., Lee, T. H., Philippe, M. & Kirschner, M. W. Cyclin activation of p34cdc2. Cell 63, 1013–1024 (1990).

49. Coudreuse, D. & Nurse, P. Driving the cell cycle with a minimal CDK control network. Nature 468, 1074–1080 (2010).

50. Shiozaki, K. & Russell, P. Cell cycle control linked to extracellular environment by MAP kinase pathway in fission yeast. 378, 739–743 (1995).

51. Petersen, J. & Hagan, I. M. Polo kinase links the stress pathway to cell cycle control and tip growth in fission yeast. Nature 435, 507–512 (2005).

52. Birot, A. et al. A second Wpl1 anti-cohesion pathway requires dephosphorylation of fission yeast kleisin Rad21 by PP4. EMBO J 36, 1364–1378 (2017).

53. Goshima, G., Iwasaki, O., Obuse, C. & Yanagida, M. The role of Ppe1/PP6 phosphatase for equal chromosome segregation in fission yeast kinetochore. EMBO J 22, 2752–2763 (2003).

54. Grallert, A. et al. A PP1-PP2A phosphatase relay controls mitotic progression. Nature 517, 94–98 (2015).

55. Rogers, S. et al. PP1 initiates the dephosphorylation of MASTL, triggering mitotic exit and bistability in human cells. J Cell Sci 129, 1340–1354 (2016).

56. Stern, B. & Nurse, P. A quantitative model for the cdc2 control of S phase and mitosis in fission yeast. Trends Genet. 12, 345–350 (1996).

57. Oikonomou, C. & Cross, F. R. Rising cyclin-CDK levels order cell cycle events. PLoS One 6, (2011).

58. McCloy, R. A. et al. Partial inhibition of Cdk1 in G2 phase overrides the SAC and decouples mitotic events. Cell Cycle 13, 1400–1412 (2014).

59. Basu, S., Patterson, J. O., Zeisner, T. U. & Nurse, P. A CDK activity buffer ensures mitotic completion. J Cell Sci 135, 1–7 (2022).

60. Moreno, S., Klar, A. & Nurse, P. Molecular genetic analysis of fission yeast Schizosaccharomyces pombe. Methods Enzymol. 194, 795–823 (1991).

61. Tanaka, N. et al. Cell Surface Galactosylation Is Essential for Nonsexual Flocculation in Schizosaccharomyces pombe. J Bacteriol 181, 1356 (1999).

62. Edelstein, A. D. et al. Advanced methods of microscope control using μManager software. J Biol Methods 1, e10 (2014).

63. Kapadia, N. et al. Processive Activity of Replicative DNA Polymerases in the Replisome of Live Eukaryotic Cells. Mol Cell 80, 114–126.e8 (2020).

64. Roberts, E. L. et al. CDK activity at the centrosome regulates the cell cycle. Cell Rep 43, (2024).

65. Colaert, N., Helsens, K., Martens, L., Vandekerckhove, J. & Gevaert, K. Improved visualization of protein consensus sequences by iceLogo. Nat Methods 6, 786–787 (2009).

66. Ge, S. X., Jung, D., Jung, D. & Yao, R. ShinyGO: A graphical gene-set enrichment tool for animals and plants. Bioinformatics 36, 2628–2629 (2020).

67. Krystkowiak, I. & Davey, N. E. SLiMSearch: A framework for proteome-wide discovery and annotation of functional modules in intrinsically disordered regions. Nucleic Acids Res 45, W464–W469 (2017).

68. Chica, N., Portantier, M., Nyquist-Andersen, M., Espada-Burriel, S. & Sandra Lopez-Aviles. Uncoupling of Mitosis and Cytokinesis Upon a Prolonged Arrest in Metaphase Is Influenced by Protein Phosphatases and Mitotic Transcription in Fission Yeast. 10, (2022).

69. Jiang, W. & Hallberg, R. L. Isolation and Characterization of par1+ and par2+: Two Schizosaccharomyces pombe Genes Encoding B′ Subunits of Protein Phosphatase 2A. Genetics 154, 1025–1038 (2000).

70. Clifford, D. M. et al. The Clp1/Cdc14 phosphatase contributes to the robustness of cytokinesis by association with anillin-related Mid1. Journal of Cell Biology 181, 79–88 (2008).

71. Mishra, M. et al. The Clp1p/Flp1p phosphatase ensures completion of cytokinesis in response to minor perturbation of the cell division machinery in Schizosaccharomyces pombe. J Cell Sci 117, 3897–3910 (2004).

72. Ohkura, H., Kinoshita, N., Miyatani, S., Toda, T. & Yanagida, M. The fission yeast dis2+ gene required for chromosome disjoining encodes one of two putative type 1 protein phosphatases. Cell 57, 997–1007 (1989).

73. Fowle, H. et al. Pp2a/b55a substrate recruitment as defined by the retinoblastoma-related protein p107. Elife 10, 1–26 (2021).

74. Hendrickx, A. et al. Docking Motif-Guided Mapping of the Interactome of Protein Phosphatase-1. Chem Biol 16, 365–371 (2009).

75. Wakula, P., Beullens, M., Ceulemans, H., Stalmans, W. & Bollen, M. Degeneracy and function of the ubiquitous RVXF motif that mediates binding to protein phosphatase-1. Journal of Biological Chemistry 278, 18817–18823 (2003).

